# DeepAffinity: Interpretable Deep Learning of Compound-Protein Affinity through Unified Recurrent and Convolutional Neural Networks

**DOI:** 10.1101/351601

**Authors:** Mostafa Karimi, Di Wu, Zhangyang Wang, Yang shen

## Abstract

**Motivation:** Drug discovery demands rapid quantification of compound-protein interaction (CPI). However, there is a lack of methods that can predict compound-protein affinity from sequences alone with high applicability, accuracy, and interpretability.

**Results:** We present a seamless integration of domain knowledges and learning-based approaches. Under novel representations of structurally-annotated protein sequences, a semi-supervised deep learning model that unifies recurrent and convolutional neural networks has been proposed to exploit both unlabeled and labeled data, for jointly encoding molecular representations and predicting affinities. Our representations and models outperform conventional options in achieving relative error in IC_50_ within 5-fold for test cases and 20-fold for protein classes not included for training. Performances for new protein classes with few labeled data are further improved by transfer learning. Furthermore, separate and joint attention mechanisms are developed and embedded to our model to add to its interpretability, as illustrated in case studies for predicting and explaining selective drug-target interactions. Lastly, alternative representations using protein sequences or compound graphs and a unified RNN/GCNN-CNN model using graph CNN (GCNN) are also explored to reveal algorithmic challenges ahead.

**Availability:** Data and source codes are available at https://github.com/Shen-Lab/DeepAffinity

**Supplementary information:** Supplementary data are available at http://shen-lab.github.io/deep-affinity-bioinf18-supp-rev.pdf.

## 1. Introduction

Drugs are often developed to target proteins that participate in many cellular processes. Among almost 900 FDA-approved drugs as of year 2016, over 80% are small-molecule compounds that act on proteins for drug effects (Santos *et al.*, 2017). Clearly, it is of critical importance to characterize compound-protein interaction for drug discovery and development, whether screening compound libraries for given protein targets to achieve desired effects or testing given compounds against possible off-target proteins to avoid undesired effects. However, experimental characterization of every possible compound-protein pair can be daunting, if not impossible, considering the enormous chemical and proteomic spaces. Computational prediction of compound-protein interaction (CPI) has therefore made much progress recently, especially for repurposing and repositioning known drugs for previously unknown but desired new targets (Keiser *et al.*, 2009; Power *et al.*, 2014) and for anticipating compound side-effects or even toxicity due to interactions with off-targets or other drugs (Chang *et al.*, 2010; Mayr *et al.*, 2016).

Structure-based methods can predict compound-protein affinity, i.e., how active or tight-binding a compound is to a protein; and their results are highly interpretable. This is enabled by evaluating energy models (Gilson and Zhou, 2007) on 3D structures of protein-compound complexes. As these structures are often unavailable, they often need to be first predicted by “docking” individual structures of proteins and compounds together before their energies can be evaluated, which tends to be a bottleneck for computational speed and accuracy (Leach *et al.*, 2006). Machine learning has been used to improve scoring accuracy based on energy features (Ain *et al.*, 2015).

More recently, deep learning has been introduced to predict compound activity or binding-affinity from 3D structures directly. Wallach et al. developed AtomNet, a deep convolutional neural network (CNN), for modeling bioactivity and chemical interactions (Wallach *et al.*, 2015). Gomes et al. (Gomes *et al.*, 2017) developed atomic convolutional neural network (ACNN) for binding affinity by generating new pooling and convolutional layers specific to atoms. Jimenez et al. (Jimenez *et al.*, 2018) also used 3D CNN with molecular representation of 3D voxels assigned to various physicochemical property channels. Besides these 3D CNN methods, Cang and Wei represented 3D structures in novel 1D topology invariants in multiple channels for CNN (Cang and Wei, 2017). These deep learning methods often improve scoring thanks to modeling long-range and multi-body atomic interactions. Nevertheless, they still rely on actual 3D structures of CPI and remain largely untested on lower-quality structures predicted from docking, which prevents large-scale applications.

Sequence-based methods overcome the limited availability of structural data and the costly need of molecular docking. Rather, they exploit rich omics-scale data of protein sequences, compound sequences (e.g. 1D binary substructure fingerprints (Wang *et al.*, 2009)) and beyond (e.g. biological networks). However, they have been restricted to classifying CPIs (Chen *et al.*, 2016) mainly into two types (binding or not) and occasionally more (e.g., binding, activating, or inhibiting (Wang and Zeng, 2013)). And more importantly, their interpretablity is rather limited due to high-level features. Earlier sequence-based machine learning methods are based on shallow models for supervised learning, such as support vector machines, logistic regression, random forest, and shallow neural networks (Cheng *et al.*, 2012; Yu *et al.*, 2012; Tabei and Yamanishi, 2013; Shi *et al.*, 2013; Cheng *et al.*, 2016). These shallow models are not lack of interpretability *per se*, but the sequence-based high-level features do not provide enough interpretability for mechanistic insights on why a compound–protein pair interacts or not.

Deep learning has been introduced to improve CPI identification from sequence data and shown to outperform shallow models. Wang and Zeng developed a method to predict three types of CPI based on restricted Boltzmann machines, a two-layer probabilistic graphical model and a type of building block for deep neural networks (Wang and Zeng, 2013). Tian et al. boosted the performance of traditional shallow-learning methods by a deep learning-based algorithm for CPI (Tian *et al.*, 2016). Wan et al. exploited feature embedding algorithm such as latent semantic algorithm (Deerwester *et al.*, 1990) and word2vec (Mikolov *et al.*, 2013) to automatically learn low-dimensional feature vectors of compounds and proteins from the corresponding large-scale unlabeled data (Wan and Zeng, 2016). Later, they trained deep learning to predict the likelihood of their interaction by exploiting the learned low-dimensional feature space. However, these deep-learning methods inherit from sequence-based methods two limitations: simplified task of predicting whether rather than how active CPIs occur as well as low interpretability due to the lack of fine-resolution structures. In addition, interpretability for deep learning models remains a challenge albeit with fast progress especially in a model-agnostic setting (Ribeiro *et al.*, 2016; Koh and Liang, 2017).

As has been reviewed, structure-based methods predict quantitative levels of CPI in a realistic setting and are highly interpretable with structural details. But their applicability is restricted by the availability of structure data, and the molecular docking step makes the bottleneck of their efficiency. Meanwhile, sequence-based methods often only predict binary outcomes of CPI in a simplified setting and are less interpretable in lack of mechanism-revealing features or representations; but they are broadly applicable with access to large-scale omics data and generally fast with no need of molecular docking.

Our goal is to, realistically, predict quantitative levels of CPIs (compound-protein affinity measured in IC_50_, *K*_*i*_, or *K*_*d*_) from sequence data alone and to balance the trade-offs of previous structure- or sequence-based methods for broad applicability, high throughput and more interpretability. From the perspective of machine learning, this is a much more challenging regression problem compared to the classification problem seen in previous sequence-based methods.

To tackle the problem, we have designed interpretable yet compact data representations and introduced a novel and interpretable deep learning framework that takes advantage of both unlabeled and labeled data. Specifically, we first have represented compound sequences in the Simplified Molecular-Input Line-Entry System (SMILES) format (Weininger, 1988) and protein sequences in novel alphabets of structural and physicochemical properties. These representations are much lower-dimensional and more informative compared to previously-adopted small-molecule substructure fingerprints or protein Pfam domains (Tian *et al.*, 2016). We then leverage the wealth of abundant unlabeled data to distill representations capturing long-term, nonlinear dependencies among residues/atoms in proteins/compounds, by pre-training bidirectional recurrent neural networks (RNNs) as part of the seq2seq auto-encoder that finds much success in modeling sequence data in natural language processing (Kalchbrenner and Blunsom, 2013). And we develop a novel deep learning model unifying RNNs and convolutional neural networks (CNNs), to be trained from end to end (Wang *et al.*, 2016b) using labeled data for task-specific representations and predictions. Furthermore, we introduce several attention mechanisms to interpret predictions by isolating main contributors of molecular fragments or their pairs, which is further exploited for predicting binding sites and origins of binding specificity. Lastly, we explore alternative representations using protein sequences or compound graphs (structural formulae), develop graph CNN (GCNN) in our unified RNN/GCNN-CNN model, and discuss remaining challenges.

The overall pipeline of our unified RNN-CNN method for semi-supervised learning (data representation, unsupervised learning, and joint supervised learning) is illustrated in Fig. 1 with details given next.

**Fig. 1.**
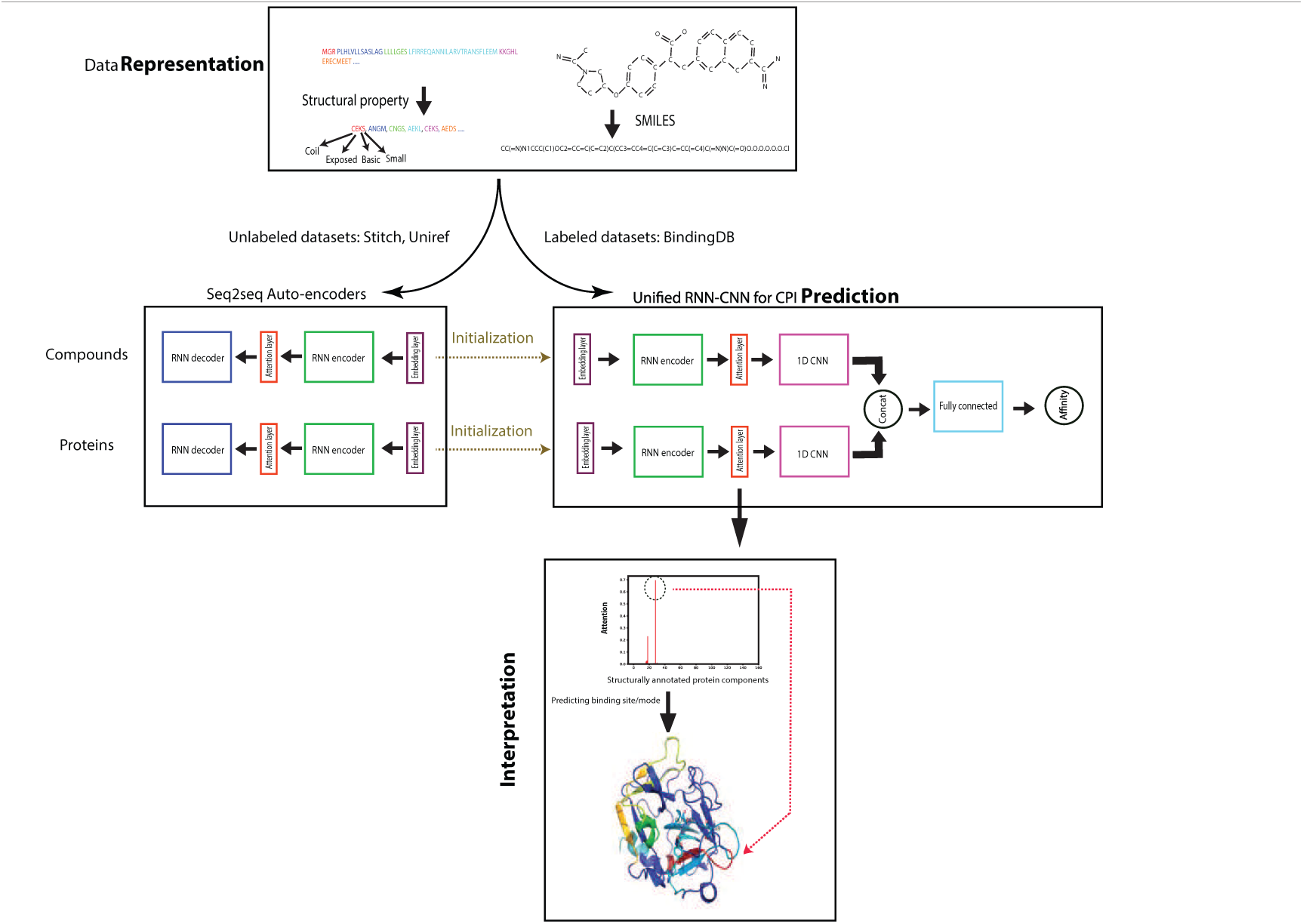
The pipeline of our unified RNN-CNN method to predict and interpret compound-protein affinity.

## 2. Materials and Methods

### 2.1 Data

We used molecular data from three public datasets: labeled compound-protein binding data from BindingDB (Liu *et al.*, 2006), compound data in the SMILES format from STITCH (Kuhn *et al.*, 2007) and protein amino-acid sequences from UniRef (Suzek *et al.*, 2014).

From 489,280 IC_50_-labeled samples collected from BindingDB, we completely excluded four classes of proteins from the training set: nuclear estrogen receptors (ER; 3,374 samples), ion channels (14,599 samples), receptor tyrosine kinases (34,318 samples), and G-protein-coupled receptors (GPCR; 60,238 samples), to test the generalizability of our framework. And we randomly split the rest into the training (263,583 samples including 10% held out for validation) and the default test set (113,168 samples) without the aforementioned four classes of protein targets. Similarly, we split a *K*_*i*_ (*K*_*d*_) labeled dataset into 101,134 (8,778) samples for training, 43,391 (3,811) for testing, 516 (4) for ERs, 8,101 (366) for ion channels, 3,355 (2,306) for tyrosine kinases, and 77,994 (2,554) for GPCRs. All labels are in logarithm forms: pIC_50_, p*K*_*i*_, and p*K*_*d*_. More details can be found in Sec. 1.1 of Supplementary Data.

For unlabeled compound data from STITCH, we randomly chose 500K samples for training and 500K samples for validation (sizes were restricted due to computing resources) and then removed those whose SMILES string lengths are above 100, resulting in 499,429 samples for training and 484,481 for validation. For unlabeled protein data from UniRef, we used all UniRef50 samples (50% sequence-identity level) less those of lengths above 1,500, resulting in 120,000 for training and 50,525 for validation.

### 2.2 Input data representation

Only 1D sequence data are assumed available. 3D structures of proteins, compounds, or their complexes are not used.

#### 2.2.1 Compound data representation

##### Baseline representation

A popular compound representation is based on 1D binary substructure fingerprints from PubChem (Wang *et al.*, 2009). Mainly, basic substructures of compounds are used as fingerprints by creating binary vectors of 881 dimensions.

##### SMILES representation

We used SMILES (Weininger, 1988) that are short ASCII strings to represent compound chemical structures based on bonds and rings between atoms. 64 symbols are used for SMILES strings in our data. 4 more special symbols are introduced for the beginning or the end of a sequence, padding (to align sequences in the same batch), or not-used ones. Therefore, we defined a compound “alphabet” of 68 “letters”. Compared to the baseline representation which uses *k*-hot encoding, canonical SMILES strings fully and uniquely determine chemical structures and are yet much more compact.

#### 2.2.2 Protein data representation

##### Baseline representation

Previously the most common protein representation for CPI classification was a 1D binary vector whose dimensions correspond to thousands of (5,523 in (Tian *et al.*, 2016)) Pfam domains (Finn *et al.*, 2014) (structural units) and 1’s are assigned based on *k*-hot encoding (Tabei and Yamanishi, 2013; Cheng *et al.*, 2016). We considered all types of Pfam entries (family, domain, motif, repeat, disorder, and coiled coil) for better coverage of structural descriptions, which leads to 16,712 entries (Pfam 31.0) as features. Protein sequences are queried in batches against Pfam using the web server HMMER (hmmscan) (Finn *et al.*, 2015) with the default gathering threshold.

##### Structural property sequence (SPS) representation

Although 3D structure data of proteins is often a luxury and their prediction remains a challenge without templates, it has been of much progress to predict protein structural properties from sequences (Cheng *et al.*, 2005; Magnan and Baldi, 2014; Wang *et al.*, 2016a). We used SSPro/ACCPro (Magnan and Baldi, 2014) to predict secondary structure class (*α*-helix, *β*-strand, and coil) and solvent accessibility (exposed or not) for each residue and group neighboring residues of the same secondary structure class into secondary structure elements (SSEs). The details and the pseudo-code for SSE are in Algorithm 1 (Supplementary Data).

Each SSE is further classified: solvent exposed if at least 30% of residues are and buried otherwise; polar, non-polar, basic or acidic based on the highest odds (for each type, occurrence frequency in the SSE is normalized by background frequency seen in all protein sequences to remove the effect from group-size difference); short if length *L* ⩽ 7, medium if 7 *< L* ⩽ 15, and long if *L >* 15. In this way, we defined 4 separate alphabets of 3, 2, 4 and 3 letters, respectively to characterize SSE category, solvent accessibility, physicochemical characteristics, and length (Table S1) and combined letters from the 4 alphabets in the order above to create 72 “words” (4-tuples) to describe SSEs. Pseudo-code for the protein representation is shown as Algorithm 2 in Supplementary Data. Considering the 4 more special symbols introduced similarly for compound SMILES strings, we flattened the 4-tuples and thus defined a protein SPS “alphabet” of 76 “letters”.

The SPS representation overcomes drawbacks of Pfam-based baseline representation: it provides higher resolution of sequence and structural details for more challenging regression tasks, more distinguishability among proteins in the same family, and more interpretability on which protein segments (SSEs here) are responsible for predicted affinity. All these are achieved with a much smaller alphabet of size 76, which leads to around 100-times more compact representation of a protein sequence than the baseline. In addition, the SPS sequences are much shorter than amino-acid sequences and prevents convergence issues when training RNN and LSTM for sequences longer than 1,000 (Li *et al.*, 2018).

### 2.3 RNN for unsupervised pre-training

We encode compound SMILES or protein SPS into representations, first by unsupervised deep learning from abundant unlabeled data. We used a recurrent neural network (RNN) model, seq2seq (Sutskever *et al.*, 2014), that has seen much success in natural language processing and was recently applied to embedding compound SMILES strings into fingerprints (Xu *et al.*, 2017). A Seq2seq model is an auto-encoder that consists of two recurrent units known as the encoder and the decoder, respectively (see the corresponding box in Fig. 1). The encoder maps an input sequence (SMILES/SPS in our case) to a fixed-dimension vector known as the thought vector. Then the decoder maps the thought vector to the target sequence (again, SMILES/SPS here). We choose gated recurrent unit (GRU) (Cho *et al.*, 2014) as our default seq2seq model and treat the thought vectors as the representations learned from the SMILES/SPS inputs. The detailed GRU configuration and advanced variants (bucketing, bidirectional GRU, and attention mechanism which provides a way to “focus” for encoders) can be found in Sec. 1.4 of Supplementary Data.

Through unsupervised pre-training, the learned representations capture nonlinear joint dependencies among protein residues or compound atoms that are far from each other in sequence. Such “long-term” dependencies are very important to CPIs since corresponding residues or atoms can be close in 3D structures and jointly contribute to intermolecular interactions.

### 2.4 Unified RNN-CNN for supervised learning

With compound and protein representations learned from the above unsupervised learning, we solve the regression problem of compound-protein affinity prediction using supervised learning. For either proteins or compounds, we append a CNN after the RNN (encoders and attention models only) that we just trained. The CNN model consists of a one-dimensional (1D) convolution layer followed by a max-pooling layer. The outputs of the two CNNs (one for proteins and the other for compounds) are concatenated and fed into two more fully connected layers.

The entire RNN-CNN pipeline is trained from end to end (Wang *et al.*, 2016b), with the pre-trained RNNs serving as warm initializations, for improved performance over two-step training. The pre-trained RNN initializations prove to be very important for the non-convex training process (Sutskever *et al.*, 2013). In comparison to such a “unified” model, we also include the “separate” RNN-CNN baseline for comparison, in which we fixed the learned RNN part and train CNN on top of its outputs.

### 2.5 Attention mechanisms in unified RNN-CNN

We have also introduced three attention mechanisms to unified RNN-CNN models. The goal is to both improve predictive performances and enable model interpretability at the level of “letters” (SSEs in proteins and atoms in compounds) and their pairs.

1. Separate attention. This default attention mechanism is applied to the compound and the protein separately so the attention learned on each side is non-specific to a compound-protein pair. However, it has the least parameters among the three mechanisms.
2. Marginalized attention. To introduce pair-specific attentions, we first use a pairwise “interaction” matrix for a pair and then marginalize it based on maximization over rows or columns for separate compound or protein attention models, which is motivated by Lu *et al.* (2016).
3. Joint attention. We have developed this novel attention model to fully explain the pairwise interactions between components (compound atoms and protein SSEs). Specifically, we use the same pairwise interaction matrix but learn to represent the pairwise space and consider attentions on pairwise interactions rather than “interfaces” on each side. Among the three attention mechanisms, joint attention provides the best interpretability albeit with the most parameters.

These attention models (for proteins, compounds, or their pairs) are jointly trained with the RNN encoder and the CNN part. Learned parameters of theirs include attention weights on all “letters” for a given string (or those on all letter-pairs for a given string-pair). Compared to that in unsupervised learning, each attention model here outputs a single vector as the input to its corresponding subsequent 1D-CNN model.

More details on unified RNN-CNN and attention mechanisms can be found in Sec. 1.5 of Supplementary Data.

## 3. Results and Discussion

### 3.1 Compound and protein representations

We compared the auto-encoding performances of our vanilla seq2seq model and 4 variants: bucketing, bi-directional GRU (“fw+bw”), attention mechanism, and attention mechanism with fw+bw, respectively, in Tables S3 and S4 (Supplementary Data). We used the common assessment metric in language models, perplexity, which is related to the entropy *H* of modeled probability distribution *P* (Perp(*P*) = 2^*H*(*P*)^ ⩾1). First, the vanilla seq2seq model had lower test-set perplexity for compound SMILES than protein SPS (7.07 versus 41.03), which echoes the fact that, compared to protein SPS strings, compound SMILES strings are defined in an alphabet of less letters (68 versus 76) and are of shorter lengths (100 versus 152), thus their RNN models are easier to learn. Second, bucketing, the most ad-hoc option among all, did not improve the results much. Third, whereas bi-directional GRUs lowered perplexity by about 2∼3.5 folds and the default attention mechanism did much more for compounds or proteins, they together achieved the best performances (perplexity being 1.0002 for compound SMILES and 1.001 for protein SPS).

Therefore, the last seq2seq variant, bidirectional GRUs with attention mechanism, is regarded the most appropriate one for learning compound/protein representations and adopted thereinafter.

### 3.2 Compound-protein affinity prediction

#### 3.2.1 Comparing novel representations to baseline ones

To assess how useful the learned/encoded protein and compound representations are for predicting compound-protein affinity, we compared the novel and baseline representations in affinity regression using the labeled datasets. The representations were compared under the same shallow machine learning models — ridge regression, lasso regression and random forest (RF).

From Table 1 we found that our novel representations learned from SMILES/SPS strings by seq2seq models outperform baseline representations of *k*-hot encoding of molecular/Pfam features. For the best performing random forest models, using 46% less training time and 24% less memory, the novel representations achieved the same performance over the default test set as the baseline ones and lowered root mean squared errors (RMSE) for two of the four generalization sets whose target protein classes (nuclear estrogen receptors / ER and ion channels) are not included in the training set. Similar improvements were observed on p*K*_*i*_, p*K*_*d*_, and pEC_50_ predictions in Tables S5–7 (Supplementary Data), respectively. These results show that learning protein and compound representations from even unlabeled datasets alone could improve their context-relevance for various labels. We also note that, unlike Pfam-based protein representations that exploit curated information only available to some proteins and their homologs, our SPS representations do not assume such information and can apply to uncharacterized proteins lacking annotated homologs.

**Table 1.** Comparing the novel representations to the baseline based on RMSE (and Pearson correlation coefficient r) of pIC_50_ shallow regression.

|  | Baseline representations |  |  | Novel representations |  |  |
| --- | --- | --- | --- | --- | --- | --- |
|  | Ridge | Lasso | RF | Ridge | Lasso | RF |
| Training | 1.16 (0.60) | 1.16 (0.60) | 0.76 (0.86) | 1.23 (0.54) | 1.22 (0.55) | **0.63** (0.91) |
| Testing | 1.16 (0.60) | 1.16 (0.60) | **0.91** (0.78) | 1.23 (0.54) | 1.22 (0.55) | **0.91** (0.78) |
| ER | 1.43 (0.30) | 1.43 (0.30) | 1.44 (0.37) | 1.46 (0.18) | 1.48 (0.18) | **1.41** (0.26) |
| Ion Channel | 1.32 (0.22) | 1.34 (0.20) | 1.30 (0.22) | 1.26 (0.23) | 1.32 (0.17) | **1.24** (0.30) |
| GPCR | **1.28** (0.22) | 1.30 (0.22) | 1.32 (0.28) | 1.34 (0.20) | 1.37 (0.17) | 1.40 (0.25) |
| Tyrosine Kinase | **1.16** (0.38) | 1.16 (0.38) | 1.18 (0.42) | 1.50 (0.11) | 1.51 (0.10) | 1.58 (0.11) |
| Time (core hours) | 3.5 | 7.4 | 1239.8 | 0.47 | 2.78 | 668.7 |
| Memory (GB) | 7.6 | 7.6 | 8.3 | 7.3 | 7.3 | 6.3 |

#### 3.2.2 Comparing shallow and deep models

Using the novel representations we next compared the performances of affinity regression between the best shallow model (random forest) and various deep models. For both separate and unified RNN-CNN models, we tested results from a single model with (hyper)parameters optimized over the training/validation set, averaging a “parameter ensemble” of 10 models derived in the last 10 epochs, and averaging a “parameter+NN” ensemble of models with varying number of neurons in the fully connected layers ((300,100), (400,200) and (600,300)) trained in the last 10 epochs. The attention mechanism used here is the default, separate attention.

From Table 2 we noticed that unified RNN-CNN models outperform both random forest and separate RNN-CNN models (the similar performances between RF and separate RNN-CNN indicated a potential to further improve RNN-CNN models with deeper models). By using a relatively small amount of labeled data (which are usually expensive and limited), protein and compound representations learned from abundant unlabeled data can be tuned to be more task-specific. We also noticed that averaging an ensemble of unified RNN-CNN models further improves the performances especially for some generalization sets of ion channels and GPCRs. As anticipated, averaging ensembles of models reduces the variance originating from network architecture and parameter optimization thus reduces expected generalization errors. Similar observations were made for p*K*_*i*_ predictions as well (Table S8 in Supplementary Data) even when their hyper-parameters were not particularly optimized and simply borrowed from pIC_50_ models. Impressively, unified RNN-CNN models without very deep architecture could predict IC_50_ values with relative errors below 10^0.7^ =5 fold (or 1.0 kcal/mol) for the test set and even around 10^1.3^ = 20 fold (or 1.8 kcal/mol) on average for protein classes not seen in the training set. Interestingly, GPCRs and ion channels had similar RMSE but more different Pearson’s *r*, which is further described by the distributions of predicted versus measured pIC_50_ values for various sets (Fig. S5 in Supplementary Data).

**Table 2.** Under novel representations learned from seq2seq, comparing random forest and variants of separate RNN-CNN and unified RNN-CNN models based on RMSE (and Pearson correlation coefficient r) for pIC_50_ prediction.

|  | RF | Separate RNN-CNN Models |  |  | Unified RNN-CNN Models |  |  |
| --- | --- | --- | --- | --- | --- | --- | --- |
|  |  | single | parameter ensemble | parameter+NN ensemble | single | parameter ensemble | parameter+NN ensemble |
| Training | 0.63 (0.91) | 0.68 (0.88) | 0.67 (0.90) | 0.68 (0.89) | 0.47 (0.94) | 0.45 (0.95) | **0.44** (0.95) |
| Testing | 0.91 (0.78) | 0.94 (0.76) | 0.92 (0.77) | 0.90 (0.79) | 0.78 (0.84) | 0.77 (0.84) | **0.73** (0.86) |
| Generalization – ER | **1.41** (0.26) | 1.45 (0.24) | 1.44 (0.26) | 1.43 (0.28) | 1.53 (0.16) | 1.52 (0.19) | 1.46 (0.30) |
| Generalization – Ion Channel | **1.24** (0.30) | 1.36 (0.18) | 1.33 (0.18) | 1.29 (0.25) | 1.34 (0.17) | 1.33 (0.18) | 1.30 (0.18) |
| Generalization – GPCR | 1.40 (0.25) | 1.44 (0.19) | 1.41 (0.20) | 1.37 (0.23) | 1.40 (0.24) | 1.40 (0.24) | **1.36** (0.30) |
| Generalization – Tyrosine Kinase | 1.58 (0.11) | 1.66 (0.09) | 1.62 (0.10) | 1.54 (0.12) | 1.24 (0.39) | 1.25 (0.38) | **1.23** (0.42) |

#### 3.2.3 Comparing attention mechanisms in prediction

To assess the predictive powers of the three attention mechanisms introduced, we compared their pIC_50_ predictions in Table 3 using the same dataset and the same unified RNN-CNN models as before. All attention mechanisms had similar performances on the training and test sets. However, as we anticipated, separate attention with the least parameters edged joint attention in generalization (especially for receptor tyrosine kinases). Meanwhile, joint attention had similar predictive performances and much better interpretability, thus will be further examined in all interpretability studies in case studies for selective drugs.

**Table 3.** Under novel representations learned from seq2seq, comparing different attention mechanisms of unified RNN-CNN models based on RMSE (and Pearson correlation coefficient r) for pIC_50_ prediction.

|  | Separate attention |  |  | Marginalized attention |  |  | Joint attention |  |  |
| --- | --- | --- | --- | --- | --- | --- | --- | --- | --- |
|  | single | parameter ensemble | parameter+NN ensemble | single | parameter ensemble | parameter+NN ensemble | single | parameter ensemble | parameter+NN ensemble |
| Training | 0.47 (0.94) | 0.45 (0.95) | 0.44 (0.95) | 0.50 (0.94) | 0.47 (0.95) | 0.42 (0.96) | 0.48 (0.94) | 0.44 (0.94) | **0.40** (0.95) |
| Testing | 0.78 (0.84) | 0.77 (0.84) | **0.73** (0.86) | 0.81 (0.83) | 0.79 (0.84) | **0.73** (0.86) | 0.84 (0.82) | 0.80 (0.83) | **0.73** (0.86) |
| Generalization – ER | 1.53 (0.16) | 1.52 (0.19) | 1.46 (0.30) | 1.69 (0.20) | 1.67 (0.20) | 1.53 (0.30) | 1.78 (0.03) | 1.68 (0.04) | **1.37** (0.23) |
| Generalization – Ion Channel | 1.34 (0.17) | 1.33 (0.18) | **1.30** (0.18) | 1.63 (0.01) | 1.64 (0.06) | 1.41 (0.13) | 1.54 (0.25) | 1.53 (0.26) | 1.42 (0.26) |
| Generalization – GPCR | 1.40 (0.24) | 1.40 (0.24) | **1.36** (0.30) | 1.59 (0.17) | 1.57 (0.18) | 1.42 (0.24) | 1.53 (0.19) | 1.53 (0.19) | 1.38 (0.25) |
| Generalization – Tyrosine Kinase | 1.24 (0.39) | 1.25 (0.38) | **1.23** (0.42) | 1.69 (0.22) | 1.62 (0.25) | 1.50 (0.32) | 2.22 (0.18) | 2.17 (0.21) | 2.04 (0.17) |

#### 3.2.4 Deep transfer learning for new classes of protein targets

Using the generalization sets, we proceed to explain and address our unified RNN-CNN models’ relatively worse performances for new classes of protein targets without any training data. We chose to analyze separate attention models with the best generalization results and first noticed that proteins in various sets have different distributions in the SPS alphabet (4-tuples). In particular, the test set, ion channels/GPCRs/tyrosine kinases, and estrogen receptors are increasingly different from the training set (measured by Jensen-Shannon distances in SPS letter or SPS length distribution) (Fig. S3 in Supplementary Data), which correlated with increasingly deteriorating performance relative to the training set (measured by the relative difference in RMSE) with a Pearson correlation coefficient of 0.68 (SPS letter distribution) or 0.96 (SPS length distribution) (Fig. S4 in Supplementary Data).

To improve the performances for new classes of proteins, we compare two strategies: re-training shallow models (random forest) from scratch based on new training data alone and “transferring” original deep models (unified parameter+NN ensemble with the default separate attention) to fit new data (see details in Supplementary Data). The reason is that new classes of targets often have few labeled data that might be adequate for re-training class-specific shallow models from scratch but not for deep models with much more parameters.

As shown in Fig. 2, deep transfer learning models increasingly improved the predictive performance compared to the original deep learning models, when increasing amount of labeled data for new protein classes are made available. The improvement was significant even with 1% training coverage for each new protein class. Notably, deep transfer learning models outperformed random forest models that were re-trained specifically for each new protein class.

**Fig. 2.**
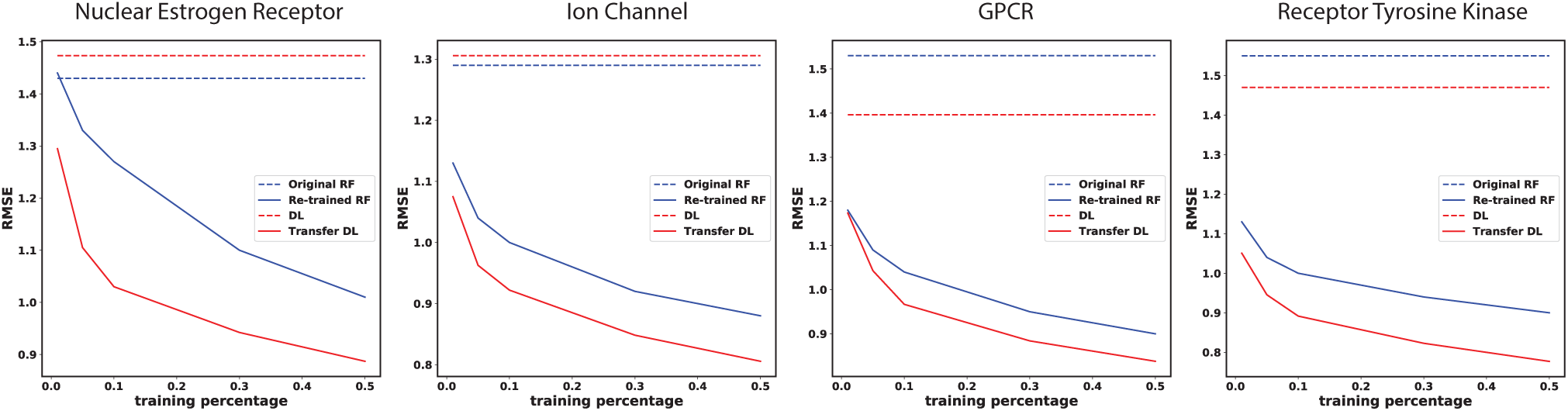
Comparing strategies to generalize predictions for four sets of new protein classes: original random forest (RF), original param.+NN ensemble of unified RNN-CNN models (DL for deep learning with the default attention), and re-trained RF or transfer DL using incremental amounts of labeled data in each set.

### 3.3 Predicting target selectivity of drugs

We went on to test how well our unified RNN-CNN models could predict certain drugs’ target selectivity, using 3 sets of drug-target interactions of increasing prediction difficulty. Our novel representations and models successfully predicted target selectivity for 6 of 7 drugs whereas baseline representations and shallow models (random forest) failed for most drugs.

#### 3.3.1 Factor Xa versus Thrombin

Thrombin and factor X (Xa) are important proteins in the blood coagulation cascade. Antithrombotics, inhibitors for such proteins, have been developed to treat cardiovascular diseases. Due to thrombin’s other significant roles in cellular functions and neurological processes, it is desirable to develop inhibitors specifically for factor Xa. DX-9065a is such a selective inhibitor (p*K*_*i*_ value being 7.39 for Xa and *<*2.70 for thrombin) (Brandstetter *et al.*, 1996).

We used the learned p*K*_*i*_ models in this study. Both proteins (thrombin and factor Xa) were included in the *K*_*i*_ training set with 2,294 and 2,331 samples, respectively, but their interactions with the compound DX-9065a were not.. Table 4 suggested that random forest correctly predicted the target selectivity (albeit with smaller than 0.5-unit difference) using baseline representations but failed to do so using novel representations. In contrast, our models with separate and joint attention mechanisms both correctly predicted the compound’s favoring Xa. Moreover, our models predicted selectivity levels being 2.4 (separate attention) and 3.9 (joint attention) in p*K*_*i*_ difference (Δp*K*_*i*_), where the joint attention model produced predictions very close to the known selectivity margin (Δp*K*_*i*_ *⩾*4.7).

**Table 4.**
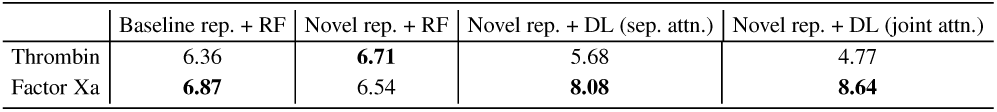
Predicted pKi values and target specificity for compound DX-9065a interacting with human factor Xa and thrombin.

#### 3.3.2 Cyclooxygenase (COX) protein family

COX protein family represents an important class of drug targets for inflammatory diseases. These enzymes responsible for prostaglandin biosynthesis include COX-1 and COX-2 in human, both of which can be inhibited by nonsteroidal anti-inflammatory drugs (NSAIDs). We chose three common NSAIDs known for human COX-1/2 selectivity: celecoxib (pIC_50_ for COX-1: 4.09; COX-2: 5.17), ibuprofen (COX-1: 4.92, COX-2: 4.10) and rofecoxib (COX-1: *<*4; COX-2: 4.6) (Luo *et al.*, 2017). This is a very challenging case for selectivity prediction because selectivity levels of all NSAIDs are close to or within 1 unit of pIC_50_.

We used the learned pIC_50_ ensemble models in this study. COX-1 and COX-2 both exist in our IC_50_ training set with 959 and 2,006 binding examples, respectively, including 2 of the 6 compound-protein pairs (ibuprofen and celecoxib with COX-1 individually).

From Table 5, we noticed that, using the baseline representations, random forest incorrectly predicted COX-1 and COX-2 to be equally favorable targets for each drug. This is because the two proteins are from the same family and their representations in Pfam domains are indistinguishable. Using the novel representations, random forest correctly predicted target selectivity for two of the three drugs (celecoxib and rofecoxib), whereas our unified RNN-CNN models (both attention mechanisms) did so for all three. Even though the selectivity levels of the NSAIDs are very challenging to predict, our models were able to predict all selectivities correctly with the caveat that few predicted differences might not be statistically significant (for instance, the 0.03-unit difference for rofecoxib using joint attention).

**Table 5.** Predicted pIC_50_ values and target specificity for three NSAIDs (CEL: celecoxib, IBU: ibuprofen and ROF: rofecoxib) interacting with human COX-1 and COX-2.

|  | Baseline rep. + RF |  |  | Novel rep. + RF |  |  | Novel rep. + DL (sep. attn.) |  |  | Novel rep. + DL (joint attn.) |  |  |
| --- | --- | --- | --- | --- | --- | --- | --- | --- | --- | --- | --- | --- |
|  | CEL | IBU | ROF | CEL | IBU | ROF | CEL | IBU | ROF | CEL | IBU | ROF |
| COX-1 | 6.06 | 5.32 | 5.71 | 6.41 | 6.12 | 6.13 | 5.11 | **6.06** | 5.67 | 5.18 | **5.94** | 6.00 |
| COX-2 | 6.06 | 5.32 | 5.71 | **6.57** | **6.19** | **6.21** | **7.60** | 5.96 | **6.51** | **7.46** | 5.62 | **6.03** |

#### 3.3.3 Protein-tyrosine phosphatase (PTP) family

Protein-tyrosine kinases and protein-tyrosine phosphatases (PTPs) are controlling reversible tyrosine phosphorylation reactions which are critical for regulating metabolic and mitogenic signal transduction processes. Selective PTP inhibitors are sought for the treatment of various diseases including cancer, autoimmunity, and diabetes. Compound 1 [2-(oxalyl-amino)-benzoic acid or OBA] and its derivatives, compounds 2 and 3 (PubChem CID: 44359299 and 90765696), are highly selective toward PTP1B rather than other proteins in the family such as PTPRA, PTPRE, PTPRC and SHP1 (Iversen *et al.*, 2000). Specifically, the p*K*_*i*_ values of OBA, compound 2, and compound 3 against PTP1B are 4.63, 4.25, and 6.69, respectively; and their p*K*_*i*_ differences to the closest PTP family protein are 0.75, 0.7, and 2.47, respectively (Iversen *et al.*, 2000).

We used the learned p*K*_*i*_ ensemble models in this study. PTP1B, PTPRA, PTPRC, PTPRE and SHP1 were included in the *K_i_* training set with 343, 33, 16, 6 and 5 samples respectively. These examples just included OBA binding to all but SHP1 and compound 2 binding to PTPRC.

Results in Table 6 showed that random forest using baseline representations cannot tell binding affinity differences within the PTP family as the proteins’ Pfam descriptions are almost indistinguishable. Using novel representations, random forest incorrectly predicted target selectivity for all 3 compounds, whereas unified RNN-CNN models with both attention mechanisms correctly did so for all but one (compound 1 – OBA). We also noticed that, although the separate attention model predicted likely insignificant selectivity levels for compounds 2 (Δp*K*_*i*_ = 0.09) and 3 (Δp*K*_*i*_ = 0.03), the joint attention model much improved the prediction of selectivity margins (Δp*K*_*i*_ = 0.58 and 0.82 for compounds 2 and 3, respectively) and their statistical significances.

**Table 6.** Predicted p*K*_*i*_ values and target specificity for three PTP1B-selective compounds interacting with five proteins in the human PTP family.

| Protein | Baseline rep. + RF |  |  | Novel rep. + RF |  |  | Novel rep. + DL (sep. attn.) |  |  | Novel rep. + DL (joint attn.) |  |  |
| --- | --- | --- | --- | --- | --- | --- | --- | --- | --- | --- | --- | --- |
|  | Comp1 | Comp2 | Comp3 | Comp1 | Comp2 | Comp3 | Comp1 | Comp2 | Comp3 | Comp1 | Comp2 | Comp3 |
| PTP1B | 4.15 | 3.87 | 5.17 | 6.70 | 6.55 | 6.71 | 3.76 | **3.84** | **3.92** | 2.84 | **4.10** | **4.04** |
| PTPRA | 4.15 | 3.87 | 5.17 | 6.29 | 6.59 | 6.27 | 2.73 | 2.90 | 3.44 | 2.39 | 2.62 | 2.12 |
| PTPRC | 4.15 | 3.87 | 5.17 | **6.86** | 6.73 | **6.87** | 3.37 | 3.25 | 3.19 | 3.36 | 3.49 | 2.97 |
| PTPRE | 4.15 | 3.87 | 5.17 | 6.79 | 6.68 | 6.81 | **3.83** | 3.75 | 3.85 | 2.75 | 2.93 | 2.61 |
| SHP1 | 4.15 | 3.87 | 5.17 | 6.71 | **6.74** | 6.73 | 3.37 | 3.38 | 3.89 | **3.42** | 3.52 | 3.22 |

### 3.4 Explaining target selectivity of drugs

After successfully predicting target selectivity for some drugs, we proceed to explain using attention scores how our deep learning models did so and what they reveal about those compound-protein interactions.

#### 3.4.1 How do the compound-protein pairs interact?

Given that SPS and SMILES strings are interpretable and attention models between RNN encoders and 1D convolution layers can report their focus, we pinpoint SSEs in proteins and atoms in compounds with high attention scores, which are potentially responsible for CPIs. To assess the idea, we chose 3 compound-protein pairs that have 3D crystal complex structures from the Protein Data Bank; and extracted residues in direct contacts with ligands (their SSEs are regarded ground truth for binding site) for each protein from ligplot diagrams provided through PDBsum (De Beer *et al.*, 2013). Based on joint attention scores *α_ij_*’s on pairs of protein SSE *i* and compound atom *j* from the single unified RNN-CNN model, we picked the top 10% (4) SSEs as predicted binding sites. Specifically, we first corrected joint attention scores to be 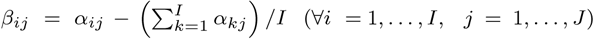 to offset the contribution of any compound atom *j* with promiscuous attentions over all protein SSEs. We then calculated the attention score *β*_*i*_ for protein SSE *i* by max-marginalization (*β*_*i*_ = max_*j*_ *β*_*ij*_). No negative *β*_*i*_ was found in this case thus no further treatment was adopted.

Table 7 shows that, compared to randomly ranking the SSEs, our approach can enrich binding site prediction by 1.7∼5.8 fold for the three CPIs. Consistent with the case of target selectivity prediction, joint attention performed better than separate attention did (Table S9). One-sided paired *t*-tests (see details in Sec. 1.7 of Supplementary Data) suggested that binding sites enjoyed higher attention scores than non-binding sites in a statistically significant way. When the strict definition of binding sites is relaxed to residues within 5Å of any heavy atom of the ligand, results were further improved with all top 10% SSEs of factor Xa being at the binding site (Table S10).

**Table 7.** Interpreting deep learning models: predicting binding sites based on joint attentions.

| Target-Drug Pair | PDB ID | Number of SSEs |  | Top 10% (4) SSEs predicted as binding site by joint attn. |  |  |  |
| --- | --- | --- | --- | --- | --- | --- | --- |
|  |  | total | binding site | # of TP | Enrichment | Best rank | P value |
| Human COX2-rofecoxib | 5KIR | 40 | 6 | 1 | 1.68 | 4 | 1.1e-2 |
| Human PTP1B-OBA | 1C85 | 34 | 5 | 1 | 1.70 | 1 | 1.1e-10 |
| Human factor Xa-DX9065 | 1FAX | 31 | 4 | 3 | 5.81 | 2 | 2.2e-16 |

We delved into the predictions for factor Xa–DX-9065a interaction in Fig. 3 (the other 2 are in Fig. S6 of Supplementary Data). Warmer colors (higher attentions) are clearly focused near the ligand. The red loops connected through a *β* strand (resi. 171–196) were correctly predicted to be at the binding site with a high rank 2, thus a true positive (TP). The SSE ranked first, a false positive, is its immediate neighbor in sequence (resi. 162-170; red helix at the bottom) and is near the ligand. In fact, as mentioned before, when the binding site definition is relaxed, all top 10% SSEs were at the binding site. Therefore, in the current unified RNN-CNN model with attention mechanism, wrong attention could be paid to sequence neighbors of ground truth; and additional information (for instance, 2D contact maps or 3D structures of proteins, if available) could be used as additional inputs to reduce false negatives.

**Fig. 3.**
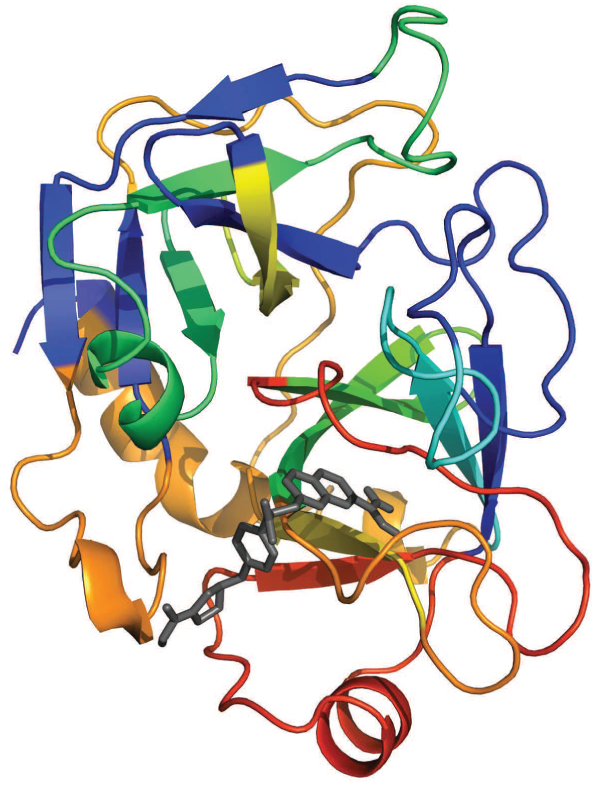
Interpreting deep learning models for factor Xa binding-site prediction based on joint attention: 3D structure of factor Xa (colored cartoons including helices, sheets, and coils) in complex with DX-9065a (black sticks) (PDB ID:1FAX) where protein SSEs are color-coded by attention scores (*β*i) where warmer colors indicate higher attentions.

We also max-marginalized *β*_*ij*_ over protein SSE *i* for *β*_*j*_ – attention score on atom *j* of the compound. Many high attention scores were observed for compound atoms (Fig. S7), which is somewhat intuitive as small-molecule compounds usually fit in protein pockets or grooves almost entirely. The top-ranked atom happened to be a nitrogen atom forming a hydrogen bond with an aspartate (Asp189) of factor Xa, although more cases need to be studied more thoroughly for a conclusion.

#### 3.4.2 How are targets selectively interacted?

To predictively explain the selectivity origin of compounds, we designed an approach to compare attention scores between pairs of CPIs and tested it using factor Xa-selective DX-9065a with known specificity origin.

For selective compounds that interact with factor Xa over thrombin, position 192 has been identified: it is a charge-neutral polar glutamine (Gln192) in Xa but a negatively-charged glutamate (Glu192) in thrombin (Huggins *et al.*, 2012). DX-9065a exploited this difference with a carboxylate group forming unfavorable electrostatic repulsion with Glu192 in thrombin but favorable hydrogen bond with Gln192 in Xa. To compare DX-9065a interacting with the two proteins, we performed amino-acid sequence alignment between the proteins and split two sequences of mis-matched SSEs (count: 31 and 38) into those of perfectly matched segments (count: 50 and 50). In the end, segment 42, where SSE 26 of Xa and SSE 31 of thrombin align, is the ground truth containing position 192 for target selectivity.

For DX-9065a interacting with either factor Xa or thrombin, we ranked the SSEs based on the attention scores from the unified RNN-CNN single model and assigned each segment the same rank as its parent SSE. Due to the different SSE counts between thrombin and factor Xa, we normalized each rank for segment *i* by the corresponding SSE count for a rank ratio *r*^*i*^. For each segment we then substracted from 1 the average of rank ratios between factor Xa and thrombin interactions so that highly attended segments in both proteins can be scored higher. Fig. 4 shows that the ground-truth segment in red was ranked the 2nd among 50 segments albeit with narrow margins over the next 3 segments.

**Fig. 4.**
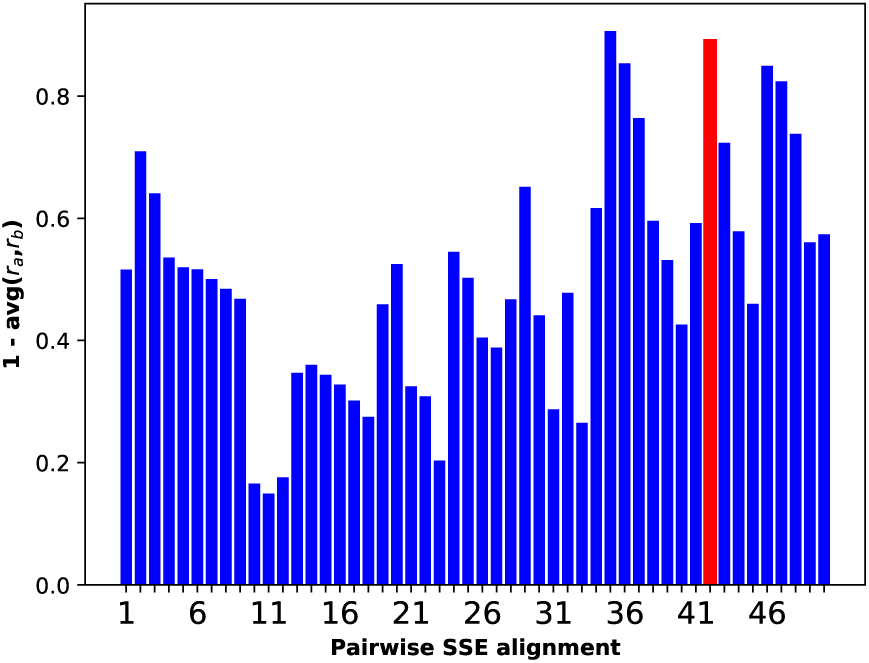
Interpreting deep learning models for factor Xa specificity based on joint attentions. Pairwise alignment of amino-acid sequences of factor Xa and thrombin decomposed both sequences into 50 segments (labeled by indices). These segments are scored by one less the average of the corrected attention rank ratios for the two compound-protein interactions. The ground truth of specificity origin is in red.

## 4. Discussion

We lastly explore alternative representations of proteins and compounds and discuss remaining challenges.

### 4.1 Protein representations using amino acid sequences

As shown earlier, our SPS representations integrate both sequence and structure information of proteins and are much more compact compared to the original amino acid sequences. That being said, there is a value to consider a protein sequence representation with the resolution of residues rather than SSEs: potentially higher-resolution precision and interpretability. We started with unsupervised learning to encode the protein sequence representation with seq2seq. More details are given in Sec. 1.8 of Supplementary Data.

**Table 8.**
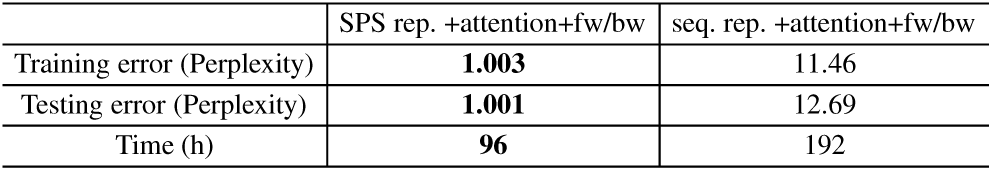
Comparing the auto-encoding performances between amino acid and SPS sequences using the best seq2seq model (bidirectional GRUs with attention mechanism).

|  | SPS rep. +attention+fw/bw | seq. rep. +attention+fw/bw |
| --- | --- | --- |
| Training error (Perplexity) | **1.003** | 11.46 |
| Testing error (Perplexity) | **1.001** | 12.69 |
| Time (h) | **96** | 192 |

Compared to SPS representations, protein sequences are 10-times longer and demanded 10-times more GRUs in seq2seq, which suggests much more expensive training. Under the limited computational budget, we trained the protein sequence seq2seq models using twice the time limit on the SPS ones. The perplexity for the test set turned out to be over 12, which is much worse than 1.001 in the SPS case (see Sec. 3.1) and deemed inadequate for subsequent (semi-)supervised learning. Learning very long sequences is challenging in general and calls for advanced architectures of sequence models.

### 4.2 Unified RNN/GCNN-CNN for protein SPS strings and compound graphs

We have chosen SMILES representations for compounds partly due to recent advancements of sequence models especially in the field of natural language processing. Meanwhile, the descriptive power of SMILES strings can have limitations. For instance, some syntactically invalid SMILES strings can still correspond to valid chemical structures. Therefore, we also explore chemical formulae (2D graphs) for compound representation.

We replaced RNN layers for compound sequences with graph CNN (GCNN) in our unified model (separate attention) and kept the rest of the architecture. This new architecture is named unified RNN/GCNN-CNN. The GCNN part is adopting a very recently-developed method (Gao *et al.*, 2018) for compound-protein interactions. More details can be found in Sec. 1.9 of Supplementary Data.

Results in Table 9 indicate that the unified RNN/GCNN-CNN model using compound graphs did not outperform the unified RNN-CNN model using compound SMILES in RMSE and did a lot worse in Pearson’s correlation coefficient. These results did not show the superiority of SMILES versus graphs for compound representations *per se*. Rather, they show that graph models need new architectures and further developments to address the challenge. We note recent advancements in deep graph models (Gilmer *et al.*, 2017; Coley *et al.*, 2017; Jin *et al.*, 2018).

**Table 9.** Comparing unified RNN-CNN (SMILES strings for compound representation) and unified RNN/GCNN-CNN (graphs for compound representation) based on RMSE (and Pearson’s correlation coefficient) for pIC_50_ prediction.

|  | SMILES rep. |  |  | Graph rep. |  |  |
| --- | --- | --- | --- | --- | --- | --- |
|  | single | parameter ensemble | parameter+NN ensemble | single | parameter ensemble | parameter+NN ensemble |
| Training | 0.47 (0.94) | 0.45 (0.95) | **0.44** (0.95) | 0.55 (0.92) | 0.54 (0.92) | 0.55 (0.92) |
| Testing | 0.78 (0.84) | 0.77 (0.84) | **0.73** (0.86) | 1.50 (0.35) | 1.50 (0.35) | 1.34 (0.45) |
| Generalization - ER | 1.53 (0.16) | 1.52 (0.19) | **1.46** (0.30) | 1.68 (0.05) | 1.67 (0.03) | 1.67 (0.07) |
| Generalization - Ion Channel | 1.34 (0.17) | 1.33 (0.18) | **1.30** (0.18) | 1.43 (0.10) | 1.41 (0.13) | 1.35 (0.12) |
| Generalization - GPCR | 1.40 (0.24) | 1.40 (0.24) | **1.36** (0.30) | 1.63 (0.04) | 1.61 (0.04) | 1.49 (0.07) |
| Generalization - Tyrosine Kinase | 1.24 (0.39) | 1.25 (0.38) | **1.23** (0.42) | 1.74 (0.01) | 1.71 (0.03) | 1.70 (0.03) |

## 5. Conclusion

We have developed accurate and interpretable deep learning models for predicting compound-protein affinity using only compound identities and protein sequences. By taking advantage of massive unlabeled compound and protein data besides labeled data in semi-supervised learning, we have jointly trained unified RNN-CNN models for learning context- and task-specific protein/compound representations and predicting compound-protein affinity. These models outperform baseline machine-learning models. And impressively, they achieve the relative error of IC_50_ within 5-fold for a comprehensive test set and even that within 10-fold for generalization sets of protein classes unknown to the training set. Deeper models would further improve the results. Moreover, for the generalization sets, we have devised transfer-learning strategies to significantly improve model performance using as few as 40 labeled samples.

Compared to conventional compound or protein representations using molecular descriptors or Pfam domains, the encoded representations learned from novel structurally-annotated SPS sequences and SMILES strings improve both predictive power and training efficiency for various machine learning models. Given the novel representations with better interpretability, we have included attention mechanism in the unified RNN-CNN models to quantify how much each part of proteins or compounds are focused while the models are making the specific prediction for each compound-protein pair.

When applied to case studies on drugs of known target-selectivity, our models have successfully predicted target selectivity in all cases whereas conventional compound/protein representations and machine learning models have failed some. Furthermore, our analyses on attention weights have shown promising results for predicting protein binding sites as well as the origins of binding selectivity, thus calling for further method development for better interpretability.

For protein representation, we have chosen SSE as the resolution for interpretability due to the known sequence-size limitation of RNN models (Li *et al.*, 2018). One can easily increase the resolution to residue-level by simply feeding to our models amino-acid sequences (preferentially of length below 1,000) instead of SPS sequences, but needs to be aware of the much increased computational burden and much worse convergence when training RNNs. For compound representation, we have started with 1D SMILES strings and have also explored 2D graph representations using graph CNN (GCNN). Although the resulting unified RNN/GCNN-CNN model did not improve against unified RNN-CNN, graphs are more descriptive for compounds and more developments in graph models are needed to address remaining challenges.

## Supporting information

## Acknowledgments

This project is in part supported by the National Institute of General Medical Sciences of the National Institutes of Health (R35GM124952 to YS) and the Defense Advanced Research Projects Agency (FA8750-18-2-0027 to ZW). Part of the computing time is provided by the Texas A&M High Performance Research Computing.

## References

Ain, Q. U., Aleksandrova, A., Roessler, F. D., and Ballester, P. J. (2015). Machinelearning scoring functions to improve structure-based binding affinity prediction and virtual screening. Wiley Interdiscip Rev Comput Mol Sci, 5(6), 405–424.

Brandstetter, H., Kühne, A., Bode, W., Huber, R., von der Saal, W., Wirthensohn, K., and Engh, R. A. (1996). X-ray structure of active site-inhibited clotting factor xa implications for drug design and substrate recognition. Journal of Biological Chemistry, 271(47), 29988–29992.

Cang, Z. and Wei, G. W. (2017). TopologyNet: Topology based deep convolutional and multi-task neural networks for biomolecular property predictions. PLoS Comput. Biol., 13(7), e1005690.

Chang, R. L., Xie, L., Xie, L., Bourne, P. E., and Palsson, B. P. (2010). Drug off-target effects predicted using structural analysis in the context of a metabolic network model. PLoS Comput. Biol., 6(9), e1000938.

Chen, X., Yan, C. C., Zhang, X., Zhang, X., Dai, F., Yin, J., and Zhang, Y. (2016). Drug-target interaction prediction: databases, web servers and computational models. Brief. Bioinformatics, 17(4), 696–712.

Cheng, F., Zhou, Y., Li, J., Li, W., Liu, G., and Tang, Y. (2012). Prediction of chemical–protein interactions: multitarget-qsar versus computational chemogenomic methods. Molecular BioSystems, 8(9), 2373–2384.

Cheng, J., Randall, A. Z., Sweredoski, M. J., and Baldi, P. (2005). Scratch: a protein structure and structural feature prediction server. Nucleic acids research, 33(suppl_2), W72–W76.

Cheng, Z., Zhou, S., Wang, Y., Liu, H., Guan, J., and Chen, Y.-P. P. (2016). Effectively identifying compound-protein interactions by learning from positive and unlabeled examples. IEEE/ACM transactions on computational biology and bioinformatics.

Cho, K., Van Merriënboer, B., Bahdanau, D., and Bengio, Y. (2014). On the properties of neural machine translation: Encoder-decoder approaches. arXiv preprint arXiv:1409.1259.

Coley, C. W., Barzilay, R., Green, W. H., Jaakkola, T. S., and Jensen, K. F. (2017). Convolutional Embedding of Attributed Molecular Graphs for Physical Property Prediction. J Chem Inf Model, 57(8), 1757–1772.

De Beer, T. A., Berka, K., Thornton, J. M., and Laskowski, R. A. (2013). Pdbsum additions. Nucleic acids research, 42(D1), D292–D296.

Deerwester, S., Dumais, S. T., Furnas, G. W., Landauer, T. K., and Harshman, R. (1990). Indexing by latent semantic analysis. Journal of the American society for information science, 41(6), 391.

Finn, R. D., Bateman, A., Clements, J., Coggill, P., Eberhardt, R. Y., Eddy, S. R., Heger, A., Hetherington, K., Holm, L., Mistry, J., Sonnhammer, E. L. L., Tate, J., and Punta, M. (2014). Pfam: the protein families database. Nucleic Acids Research, 42(D1), D222–D230.

Finn, R. D., Clements, J., Arndt, W., Miller, B. L., Wheeler, T. J., Schreiber, F., Bateman, A., and Eddy, S. R. (2015). Hmmer web server: 2015 update. Nucleic acids research, 43(W1), W30–W38.

Gao, K. Y., Fokoue, A., Luo, H., Iyengar, A., Dey, S., and Zhang, P. (2018). Interpretable drug target prediction using deep neural representation. In IJCAI, pages 3371–3377.

Gilmer, J., Schoenholz, S. S., Riley, P. F., Vinyals, O., and Dahl, G. E. (2017). Neural message passing for quantum chemistry. CoRR, abs/1704.01212.

Gilson, M. K. and Zhou, H.-X. (2007). Calculation of protein-ligand binding affinities. Annual review of biophysics and biomolecular structure, 36.

Gomes, J., Ramsundar, B., Feinberg, E. N., and Pande, V. S. (2017). Atomic convolutional networks for predicting protein-ligand binding affinity. arXiv preprint arXiv:1703.10603.

Huggins, D. J., Sherman, W., and Tidor, B. (2012). Rational approaches to improving selectivity in drug design. Journal of medicinal chemistry, 55(4), 1424–1444.

Iversen, L. F., Andersen, H. S., Branner, S., Mortensen, S. B., Peters, G. H., Norris, K., Olsen, O. H., Jeppesen, C. B., Lundt, B. F., Ripka, W., et al. (2000). Structure-based design of a low molecular weight, nonphosphorus, nonpeptide, and highly selective inhibitor of protein-tyrosine phosphatase 1b. Journal of Biological Chemistry, 275(14), 10300–10307.

Jimenez, J., Skalic, M., Martinez-Rosell, G., and De Fabritiis, G. (2018). KDEEP: Protein-Ligand Absolute Binding Affinity Prediction via 3D-Convolutional Neural Networks. J Chem Inf Model, 58(2), 287–296.

Jin, W., Barzilay, R., and Jaakkola, T. S. (2018). Junction tree variational autoencoder for molecular graph generation. CoRR, abs/1802.04364.

Kalchbrenner, N. and Blunsom, P. (2013). Recurrent continuous translation models. In EMNLP, volume 3, page 413.

Keiser, M. J., Setola, V., Irwin, J. J., Laggner, C., Abbas, A., Hufeisen, S. J., Jensen, N. H., Kuijer, M. B., Matos, R. C., Tran, T. B., et al. (2009). Predicting new molecular targets for known drugs. Nature, 462(7270), 175.

Koh, P. W. and Liang, P. (2017). Understanding black-box predictions via influence functions. In D. Precup and Y. W. Teh, editors, Proceedings of the 34th International Conference on Machine Learning, volume 70 of Proceedings of Machine Learning Research, pages 1885–1894, International Convention Centre, Sydney, Australia. PMLR.

Kuhn, M., von Mering, C., Campillos, M., Jensen, L. J., and Bork, P. (2007). Stitch: interaction networks of chemicals and proteins. Nucleic acids research, 36(suppl_1), D684–D688.

Leach, A. R., Shoichet, B. K., and Peishoff, C. E. (2006). Prediction of proteinligand interactions. Docking and scoring: successes and gaps. J. Med. Chem., 49(20), 5851–5855.

Li, S., Li, W., Cook, C., Zhu, C., and Gao, Y. (2018). Independently recurrent neural network (indrnn): Building A longer and deeper RNN. CoRR, abs/1803.04831.

Liu, T., Lin, Y., Wen, X., Jorissen, R. N., and Gilson, M. K. (2006). Bindingdb: a web-accessible database of experimentally determined protein–ligand binding affinities. Nucleic acids research, 35(Suppl_1), D198–D201.

Lu, J., Yang, J., Batra, D., and Parikh, D. (2016). Hierarchical question-image co-attention for visual question answering. In Advances In Neural Information Processing Systems, pages 289–297.

Luo, Y., Zhao, X., Zhou, J., Yang, J., Zhang, Y., Kuang, W., Peng, J., Chen, L., and Zeng, J. (2017). A network integration approach for drugtarget interaction prediction and computational drug repositioning from heterogeneous information. Nature communications, 8(1), 573.

Magnan, C. N. and Baldi, P. (2014). Sspro/accpro 5: almost perfect prediction of protein secondary structure and relative solvent accessibility using profiles, machine learning and structural similarity. Bioinformatics, 30(18), 2592–2597.

Mayr, A., Klambauer, G., Unterthiner, T., and Hochreiter, S. (2016). Deeptox: Toxicity prediction using deep learning. Frontiers in Environmental Science, 3, 80.

Mikolov, T., Chen, K., Corrado, G., and Dean, J. (2013). Efficient estimation of word representations in vector space. arXiv preprint arXiv:1301.3781.

Power, A., Berger, A. C., and Ginsburg, G. S. (2014). Genomics-enabled drug repositioning and repurposing: insights from an IOM Roundtable activity. JAMA, 311(20), 2063–2064.

Ribeiro, M. T., Singh, S., and Guestrin, C. (2016). “why should i trust you?*”: Explaining the predictions of any classifier. In Proceedings of the 22Nd ACM SIGKDD International Conference on Knowledge Discovery and Data Mining, KDD’ 16, pages 1135–1144, New York, NY, USA. ACM.

Santos, R., Ursu, O., Gaulton, A., Bento, A. P., Donadi, R. S., Bologa, C. G., Karlsson, A., Al-Lazikani, B., Hersey, A., Oprea, T. I., and Overington, J. P. (2017). A comprehensive map of molecular drug targets. Nat Rev Drug Discov, 16(1), 19–34.

Shi, Y., Zhang, X., Liao, X., Lin, G., and Schuurmans, D. (2013). Protein-chemical interaction prediction via kernelized sparse learning svm. In Pacific Symposium on Biocomputing, pages 41–52.

Sutskever, I., Martens, J., Dahl, G., and Hinton, G. (2013). On the importance of initialization and momentum in deep learning. In International conference on machine learning, pages 1139–1147.

Sutskever, I., Vinyals, O., and Le, Q. V. (2014). Sequence to sequence learning with neural networks. In Advances in neural information processing systems, pages 3104–3112.

Suzek, B. E., Wang, Y., Huang, H., McGarvey, P. B., Wu, C. H., and Consortium, U. (2014). Uniref clusters: a comprehensive and scalable alternative for improving sequence similarity searches. Bioinformatics, 31(6), 926–932.

Tabei, Y. and Yamanishi, Y. (2013). Scalable prediction of compound-protein interactions using minwise hashing. BMC systems biology, 7(6), S3.

Tian, K., Shao, M., Wang, Y., Guan, J., and Zhou, S. (2016). Boosting compound-protein interaction prediction by deep learning. Methods, 110, 64–72.

Wallach, I., Dzamba, M., and Heifets, A. (2015). Atomnet: a deep convolutional neural network for bioactivity prediction in structure-based drug discovery. arXiv preprint arXiv:1510.02855.

Wan, F. and Zeng, J. (2016). Deep learning with feature embedding for compound-protein interaction prediction. bioRxiv, page 086033.

Wang, S., Li, W., Liu, S., and Xu, J. (2016a). Raptorx-property: a web server for protein structure property prediction. Nucleic Acids Research, 44(W1), W430–W435.

Wang, Y. and Zeng, J. (2013). Predicting drug-target interactions using restricted Boltzmann machines. Bioinformatics, 29(13), i126–134.

Wang, Y., Xiao, J., Suzek, T. O., Zhang, J., Wang, J., and Bryant, S. H. (2009). Pubchem: a public information system for analyzing bioactivities of small molecules. Nucleic acids research, 37(Suppl_2), W623–W633.

Wang, Z., Chang, S., Yang, Y., Liu, D., and Huang, T. S. (2016b). Studying very low resolution recognition using deep networks. In Proceedings of the IEEE Conference on Computer Vision and Pattern Recognition, pages 4792–4800.

Weininger, D. (1988). Smiles, a chemical language and information system. 1. introduction to methodology and encoding rules. Journal of chemical information and computer sciences, 28(1), 31–36.

Xu, Z., Wang, S., Zhu, F., and Huang, J. (2017). Seq2seq fingerprint: An unsupervised deep molecular embedding for drug discovery. In Proceedings of the 8th ACM International Conference on Bioinformatics, Computational Biology, and Health Informatics, pages 285–294. ACM.

Yu, H., Chen, J., Xu, X., Li, Y., Zhao, H., Fang, Y., Li, X., Zhou, W., Wang, W., and Wang, Y. (2012). A systematic prediction of multiple drug-target interactions from chemical, genomic, and pharmacological data. PloS one, 7(5), e37608.

