## Supplementary material for "DeepAffinity: Interpretable Deep Learning of Compound-Protein Affinity through Unified Recurrent and Convolutional Neural Networks"

#### 1 Materials and Methods

##### 1.1 Data curation

For the labeled data, we chose the half maximal inhibitory concentration ($IC_{50}$) as the main measure of binding affinity and extracted samples with known $IC_{50}$ values from BindingDB (Liu *et al.*, 2006). We used the measurements along with amino-acid sequences of proteins and 2D structures of compounds provided by BindingDB in the file "BindingDB_All_2018m8.tsv.zip". Since compound SMILES strings are not always available and not necessarily in the canonical form, we retrieved these strings from PubChem (Kim *et al.*, 2016) by cross-referencing their PubChem CIDs (compound IDs). As 95% of compounds and proteins with known $IC_{50}$ values are of lengths up to 100 (SMILES characters) and 1500 (amino acids), respectively, we removed samples outside of the length ranges. And we also removed samples with incomplete information to determine protein sequences (such as those containing the letter "X", lower-case letters, or arabic numerals) or with $IC_{50}$ as a range instead of exact value (unless it is worse than a weak $10^{-2}$M (10mM) or better than a tight $10^{-11}$M (0.01nM)). Moreover, for each protein-compound pairs with multiple measurements, we removed measurements outside aforementioned ranges and then used the geometric mean of the rest. In the end, we collected 489,280 $IC_{50}$ samples from BindingDB.

The collected $IC_{50}$ data are split into training, testing, and 4 generalization sets. We first withheld 4 generalization sets of all the 3,374 nuclear estrogen receptors (ER $\alpha$ and $\beta$), 14,599 ion channels, 34,318 receptor tyrosine kinases, and 60,238 G-protein coupled receptors (GPCR), respectively; and then split the remaining into the training (263,583) and the testing (113,168) sets. To determine each of the 4 generalization sets (protein classes), we compared each protein's Gene Ontology (GO) molecular function terms to a set of target GO terms. Specifically, using the QuickGO REST API, we identified for each of the 4 protein classes a target GO molecular function term (ER: GO:0030284, ion channel: GO:0005216, receptor tyrosine kinase: GO:0004713, and GPCR: GO:0004930) and its molecular-function descendants. And using UnitProt ID as a protein identifier, we retrieved each protein's molecular function GO terms by cross-referencing its UniProt entry html. If any GO term of a protein belongs to a target GO set, we will classify the protein to the corresponding protein class. The only exception applied to ER. Since some proteins contained target GO terms for both ER and GPCR, we only classified proteins with the ESR1 and ESR2 genes (again retrieved from their UniProt entries' html files) to nuclear estrogen receptor (ER).

We used $IC_{50}$ in its logarithm form ($pIC_{50} = -\log_{10}IC_{50}$) as the target label for regression. Prior to taking the logarithm, we truncated $IC_{50}$ to the range of [$10^{-11}$M, $10^{-2}$M] considering the sensitive ranges of experimental methods for $IC_{50}$ measurement. In other words, measurements stronger than $10^{-11}$ or weaker than $10^{-2}$ would be regarded $10^{-11}$ or $10^{-2}$, respectively.

We also used $K_i$ and $K_d$ values (again, in their logarithm forms) from BindingDB as measures of binding affinity. These data were curated similarly as the $IC_{50}$ data. In total, we split a curated $K_i$ ($K_d$) dataset into 101,134 (8,778) samples for training, 43,391 (3,811) for testing, 516 (4) for ER, 8,101 (366) for ion channels, 3,355 (2,306) for receptor tyrosine kinases, and 77,994 (2,554) for GPCRs. Furthermore, we curated $EC_{50}$ data from BindingDB similarly and split them into 26,413 samples for training, 11,483 for testing, 961 for ER, 1,691 for ion channels, 292 for receptor tyrosine kinases, and 21,720 for GPCRs. Due to their relatively small sizes, $K_d$ and $EC_{50}$ data were only used in shallow regression to compare the baseline and novel representations.

#### 1.2 Protein representation

We designed an algorithm to first generate secondary structure elements (SSEs) from predicted secondary structure classes based on two rules: the minimum size of $\alpha$- or $\beta$ SSEs is 5 and that for coil SSEs is 3; a group of less size is merged to neighboring SSEs that satisfy the minimum size, by sliding a window of size 7 down the sequence to "smooth" it. The rules are adopted to address uncertain or "orphan" groups of few members. More details can be found in Algorithm 1.

Each SSE is classified based on secondary structure category, solvent accessibility, physicochemical properties, and residue length (Table S1). Specifically, an SSE is solvent exposed if at least 30% of residues are and buried otherwise; polar, non-polar, basic or acidic based on the highest odds (for each type, occurrence frequency in the SSE is normalized by background frequency seen in all protein sequences to remove the effect from group-size difference, See Table S2); short if length $L \leqslant 7$, medium if $7 < L \leqslant 15$, and long if $L > 15$. More details can be found in Algorithm 2.

| Secondary Structure | | | Solvent Exposure | | Property | | | | Length | | |
|---|---|---|---|---|---|---|---|---|---|---|---|
| Alpha | Beta | Coil | Not Exposed | Exposed | Non-polar | Polar | Acidic | Basic | Short | Medium | Long |
| A | B | C | N | E | G | T | D | K | S | M | L |

Table S1: 4-tuple of letters in protein structural property sequence (SPS) for creating words.

**Algorithm 1** "Smoothing" Secondary Structure Element Sequence

```
Input: Secondary Structures over a Protein Sequence
Output: Smoothed Secondary Structure Element Sequence
Initialize: MeaningfulMinLength as 3 for Coil and 5 for Sheet and Helix
Initialize: MaxGroupLength as 7
for Each line in Secondary Structure do
  Initialize FinalSeq as Empty
  for Each letter in line do
    if Current = Next then
      Store Current in S
      Mark the length of continue same sequence as L
    else if L > MeaningfulMinLength then
      Smooth all letters in S to last letter
      Store S to FinalSeq and clear S
      Continue
    else if Length of S < MaxGroupLength then
      Store to S
      Continue
    else
      Smooth all letters to the most number letter in S
      Store S to FinalSeq and Clear S
      Continue
    end if
  end for
  return FinalSeq
  Clear FinalSeq
end for
```

| Groups | Amino-acid members | # of members | Background frequency |
|---|---|---|---|
| Nonpolar | Gly, Ala, Val, Leu, Ile, Met, Phe, Pro, Trp | 9 | 0.505 |
| Polar | Ser, Thr, Tyr, Cys, Gln, Asn | 6 | 0.253 |
| Acidic | Asp, Glu | 2 | 0.111 |
| Basic | Lys, Arg, His | 3 | 0.131 |

Table S2: Nonpolar, polar, basic and acidic amino acids and their frequencies.

In this way, we defined 4 "alphabets" of 3, 2, 4 and 3 "letters", respectively to characterize categories in SSE, solvent accessibility, physicochemical characteristics, and length (Table S2) and combined letters from the 4 alphabets in the order above to create 72 "words" (4-tuples) to describe SSEs. For example, the word "AEKM" means an $\alpha$-type SSE that is solvent-exposed, overly basic, and of medium length. Lastly, we "flattened" the 4-tuples into a protein alphabet of 76 letters (72 plus 4 special symbols such as beginning, ending, padding, and not-used ones).

**Algorithm 2** Secondary Structure Element (SSE) Sequence to Structure Property Sequence (SPS)

```
Input: Smoothed Secondary Structure and Exposedness Protein Sequence
Output: Structural Property Sequence (SPS).
Initialize: Exposedness threshold eThres
Initialize: Polarity threshold pThres based on the average of each types
for Each ss in Secondary Structure Element Sequences do
  Initialize FinalSeq as Empty
  Read next Exposedness Protein Sequence to acc
  Read next Protein Sequence to protein
  for Each continuous same letter in ss do
    Store corresponding letter to G
    if the number of 'e' in ss > eThres then
      Store 'E' to G
    else
      Store 'N' to G
    end if
    if percentage of one type polarity in protein > pThres then
      Store corresponding letter to G
    end if
    if Length of group ≤ 7 then
      Store 'S' letter to G
    else if Length of group ≤ 15 then
      Store 'M' letter to G
    else if Length of group > 15 then
      Store 'L' letter to G
    end if
    Store G to FinalSeq
    Clear G
  end for
  return FinalSeq
  Clear FinalSeq
end for
```

##### 1.3 Shallow machine learning models

Shallow machine learning models — ridge, lasso, and random forest (RF) — were used to compare learned protein/compound representations with baseline ones and implemented using scikit-learn (Pedregosa *et al.*, 2011). Hyper parameters include regularization constant $\alpha \in \{10^{-4}, 10^{-3}, \ldots, 10^{4}\}$ for lasso and ridge regression as well as the number of trees $\in \{50, 100, 200, 500\}$, the minimum sample per leaf $\in \{10, 50, 100\}$ and the maximum number of features for each tree $\in \{n, \sqrt{n}, \log_2 n\}$ for random forest where $n$ denotes the number of features. These hyper parameters were optimized using 10-fold cross validation.

##### 1.4 RNN for Unsupervised Learning

###### 1.4.1 Introduction to GRU

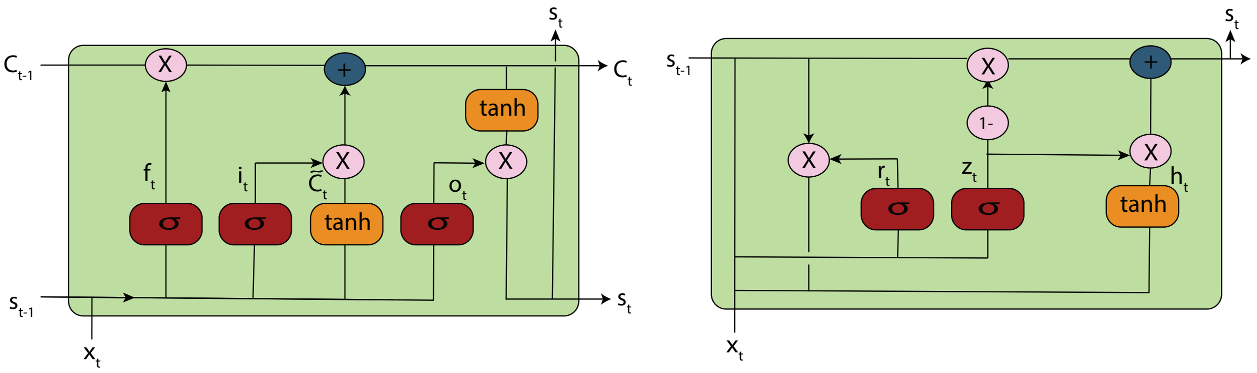


Figure S1: LSTM and GRU

As shown in Fig. S1, an LSTM unit consists of three "gates" — input ($i_t$), forget ($f_t$) and output ($o_t$) gates as well as two states — cell (also memory, $C_t$) and hidden states (also output, $s_t$). LSTM will iterate over the input sequence $(x_1, \cdots, x_t, \cdots, x_T)$ and calculate the output sequence of $(s_1, \cdots, s_t, \cdots, s_T)$ through:

$$
\begin{aligned}
f_t &= \sigma(W_f \times [s_{t-1}, x_t] + b_f) \\
i_t &= \sigma(W_i \times [s_{t-1}, x_t] + b_i) \\
\tilde{C}_t &= \tanh(W_c \times [s_{t-1}, x_t] + b_c) \\
C_t &= f_t \times C_{t-1} + i_t \times \tilde{C}_t \\
o_t &= \sigma(W_o \times [s_{t-1}, x_t] + b_o) \\
s_t &= o_t \times \tanh(C_t)
\end{aligned}
\tag{1}
$$

GRU further simplifies LSTM without sacrificing the performance much. With much less parameters, it demands lighter computational time. A GRU unit consists of two gates — reset ($r_t$) and update ($Z_t$) gates as well as one state — hidden state ($s_t$). It calculates the output sequence of $(s_1, \cdots, s_t, \cdots, s_T)$ over the input sequence $(x_1, \cdots, x_t, \cdots, x_T)$ through:

$$
\begin{aligned}
z_t &= \sigma(W_z \times [s_{t-1}, x_t] + b_z) \\
r_t &= \sigma(W_r \times [s_{t-1}, x_t] + b_r) \\
h_t &= \tanh(W_h \times [r_t \times s_{t-1}, x_t] + b_h) \\
s_t &= (1 - z_t) \times s_{t-1} + z_t \times h_t
\end{aligned}
\tag{2}
$$

##### 1.4.2 GRU implementation

Our alphabets include 68 and 76 letters (including 4 special symbols such as padding in either alphabet) for compound SMILES and protein SPS strings, respectively. Based on the statistics of 95% CPIs in BindingDB, we set the maximum lengths of SMILES and SPS strings to be 100 and 152, respectively. Accordingly, we used 2 layers of GRU with both the latent dimension and the embedding layer (discrete letter to continuous vector) dimension being 128 for compounds and 256 for proteins. We used an initial learning rate of 0.5 with a decay rate of 0.99, a dropout rate of 0.2, and a batch size of 64. We also clipped gradients by their global norms. All neural network models for supervised learning were implemented based on TensorFlow (Abadi *et al.*, 2016) and TFLearn (Tang, 2016).

We further test four more seq2seq variants under the combination of the following options:

- **Bucketing** (Khomenko *et al.*, 2016) as an optimization trick to put sequences of similar lengths in the same bucket and padded accordingly during training. We used bucketing groups of {(30,30), (60,60), (90,90), (120,120), (152,152)} for SPS and {(20,20), (40,40), (60,60), (80,80), (100,100)} for SMILES.
- **Bidirectional GRU** that shares parameters to capture both forward and backward dependencies (Schuster and Paliwal, 1997).
- **Attention** mechanism (Bahdanau *et al.*, 2014) that allows encoders to "focus" in each encoding step on selected previous time points that are deemed important to predict target sequences.

Both bidirectional RNN and attention mechanism can help address computational challenges from long input sequences.

##### 1.4.3 Attention mechanism for unsupervised learning

To address the challenge from long input sequences, the attention mechanism provides a way to "focus" for encoders. It only allows each encoding step to be affected by selected previous time points deemed important, thus saving its memory burden. Suppose that the maximum length of both the encoder and the decoder is $L$, the output of RNN (the input to the attention model) is $(s_1, \cdots, s_t, \cdots, s_L)$, the hidden state of the decoder $(h_1, \cdots, h_j, \cdots, h_L)$, and the output of attention model is $C_j$ at each time step $j$ of the decoder. The attention model is parametrized by two matrices, $U_\text{a}$ and $W_\text{a}$, and a vector $\boldsymbol{v}_\text{a}$ ('a' stands for attention) and formulated as:

$$\begin{aligned} e_{j,t} &= \boldsymbol{v}_\text{a} \tanh(U_\text{a} h_{j-1} + W_\text{a} s_t) \quad \forall j = 1, \cdots, L \quad \text{and} \quad \forall t = 1, \cdots, L \\ \alpha_{j,t} &= \frac{exp(e_{j,t})}{\sum_{k=1}^{L} exp(e_{j,k})} \quad \forall j = 1 \cdots L \quad and \quad t = 1 \cdots L \\ C_j &= \sum_{t=1}^{L} \alpha_{j,t} s_t \quad \forall j = 1 \cdots L \end{aligned} \tag{3}$$

All seq2seq models were implemented based on the seq2seq code released by Google (Britz *et al.*, 2017).

#### 1.5 Unified RNN-CNN for supervised learning

##### 1.5.1 CNN

Convolutional neural networks (CNN) (Lawrence *et al.*, 1997) have made great success in modeling image data for computer vision. For either proteins or compounds, our CNN model consisted of a one-dimensional (1D) convolution layer followed by a max-pooling layer. The outputs of these layers for proteins and compounds were concatenated then fed to two fully connected (or dense) layers.

In particular, the 1D convolution layer had 64 various filters (all have length 8 and stride 4 for proteins and length 4 and stride 2 for compounds). The max-pooling layer with a filter of length 4 was to reduce the parameters for the next layers and introduce nonlinearity to the predictor. The 2 fully connected layers are with 200 and 50 neurons, respectively, activation function of leaky RELU (Maas *et al.*, 2013), Xavier initialization (Glorot and Bengio, 2010), and a drop-out rate of 20%.

##### 1.5.2 Separate and unified RNN-CNN models

RNNs (encoders and attention models only) for unsupervised learning and CNNs for supervised learning are connected in series and trained separately or jointly (see Fig. S2).

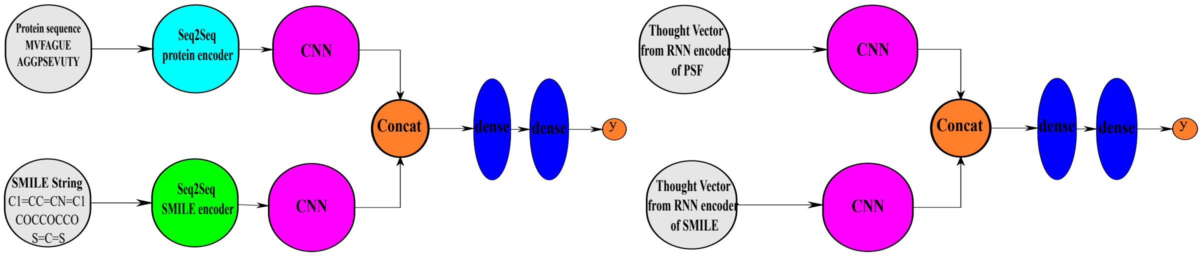


Figure S2: Separate and unified RNN-CNN models.

In separate RNN-CNN models, a seq2seq model for proteins or compounds is trained once for context-specific representations and thought vectors from its encoder are fed as the input to train a subsequent CNN model. In unified RNN-CNN models, both RNN (just encoder + attention model) and CNN models are trained together to jointly learn representations and make predictions, which makes the representations further relevant to the specific task.

Considering that training the unified model involves a non-convex optimization problem in a much higher dimensional parameter space than training the CNN model alone, we used previously-learned GRU parameters as starting points to have a "warm start" for optimization.

Specifically, we first fixed parameters of RNN encoders and trained the attention models, CNN layers and fully connected layers over the labeled training set for 100 epochs with a learning rate of $10^{-3}$. After the newly-trained layers are also warmed up (i.e., initialized), we jointly trained the unified RNN-CNN model for 200 epochs with a learning rate of $10^{-4}$.

In other words, we fine-tuned protein or compound representations learned from seq2seq previously for compound-protein affinity prediction.

##### 1.5.3 Attention mechanisms for unified RNN-CNN models

**Separate attention.** We have also introduced protein and compound attention models in supervised learning to both improve predictive performances and enable model interpretability at the level of "letters" (SSEs in proteins and atoms in compounds). In the supervised model we just have the encoder and its attention $\alpha_t$ on each letter $t$ for a given string $\mathbf{x}$ (protein or compound). And the output of the attention model, $A$, will be the input to the subsequent 1D-CNN model. Suppose that the length of protein encoder is $T$ and $(s_1, \cdots, s_t, \cdots, s_T)$ are the output of protein encoder and similarly the length of compound encoder is $D$ and $(m_1, \cdots, m_d, \cdots, m_D)$ are the output of compound encoder. We parametrize the attention model of unified model with matrix $U_\mathrm{a}$ and the vector $\boldsymbol{v}_\mathrm{a}$. Then, The attention model for the protein encoder is formulated as:

$$\begin{aligned} e_t^P &= \boldsymbol{v}_\mathrm{a}^P \tanh(W_\mathrm{a}^P s_t) \quad \forall t = 1, \cdots, T \\ \alpha_t^P &= \frac{exp(e_t^P)}{\sum_{k=1}^T exp(e_k^P)} \quad \forall t = 1 \cdots T \\ A^P &= \sum_{t=1}^T \alpha_t^P s_t \end{aligned} \tag{4}$$

Similarly for the compound encoder:

$$\begin{aligned} e_d^C &= \boldsymbol{v}_\mathrm{a}^C \tanh(W_\mathrm{a}^C m_d) \quad \forall d = 1, \cdots, D \\ \alpha_d^C &= \frac{exp(e_d^C)}{\sum_{k=1}^D exp(e_k^C)} \quad \forall d = 1 \cdots D \\ A^C &= \sum_{d=1}^D \alpha_d^C m_d \end{aligned} \tag{5}$$

The attention weights (scores) $\alpha_t^\cdot$ suggest the importance of the $t^{\mathrm{th}}$ "letter" (secondary structure element in proteins and atom or connectivity in compounds) and thus predict the binding sites relevant to the predicted binding affinity.

**Marginalized attention.** Considering that the separate attention model does not address compound-protein pair specificity, we have exploited a co-attention mechanism similar to Lu *et al.* (2016) which has been widely used in visual question answering. We name this attention mechanism "marginalized attention" because we marginalize over rows or columns of a pair-specific interaction matrix for compound or protein attention in this pair. Specifically, a pairwise "interaction" matrix $N$ of size $T \times D$ is defined with each element as:

$$N_{td} = \tanh(s_t^T W_\mathrm{a} m_d) \quad \forall t = 1, \cdots, T, \quad \forall d = 1, \cdots, D, \tag{6}$$

where the parameter matrix $W_\mathrm{a}$ is now of size $T \times D$.
By considering max column marginalization, we have the marginalized attention model for the protein:

$$\begin{aligned} e_t^P &= \max_{d=1:D}(N_{td}) \quad \forall t = 1, \cdots, T \\ \alpha_t^P &= \frac{exp(e_t^P)}{\sum_{k=1}^T exp(e_k^P)} \quad \forall t = 1 \cdots T \\ A^P &= \sum_{t=1}^T \alpha_t^P s_t \end{aligned} \tag{7}$$

Similarly, by max row marginalization, we have the marginalized attention model for compounds:

$$\begin{aligned} e_d^C &= \max_{t=1:T}(N_{td}) \quad \forall d = 1, \cdots, D \\ \alpha_d^C &= \frac{exp(e_d^C)}{\sum_{k=1}^D exp(e_k^C)} \quad \forall d = 1 \cdots D \\ A^C &= \sum_{d=1}^D \alpha_d^C m_d \end{aligned} \tag{8}$$

**Joint attention.** We have further developed a novel "joint attention" mechanism that removes the need of marginalization and thus pays attention directly on interacting "letter" pairs rather than individual interfaces. Specifically, for the same pairwise interaction matrix $N$ of size $T \times D$ as defined in Eq. (6), we would learn the space for the joint model by another layer of neural network:

$$B_{td} = \tanh(V_{\mathrm{b}} s_t + W_{\mathrm{b}} m_d + b) \quad \forall d = 1 \cdots D, \quad \forall t = 1, \cdots, T \tag{9}$$

Then we can derive the attention score $\alpha_{td}$ for each $(t, d)$ pair and the final output $A$ as:

$$\begin{aligned} \alpha_{td} &= \frac{exp(N_{td})}{\sum_{k=1}^{T} \sum_{k'=1}^{D} exp(N_{kk'})} \quad \forall d = 1 \cdots D, \quad \forall t = 1, \cdots, T \\ A &= \sum_{t=1}^{T} \sum_{d=1}^{D} \alpha_{td} B_{td} \end{aligned} \tag{10}$$

Throughout the study, we disregarded attention scores on special symbols and re-normalized the rest. For compound SMILES, we further disregarded SMILES symbols that are not English letters when interpreting compound attentions on atoms, which awaits to be improved.

All neural network models for supervised learning were implemented based on TensorFlow (Abadi *et al.*, 2016) and TFLearn (Tang, 2016).

#### 1.6 Deep transfer learning

For deep transfer learning, we fine-tuned the first embedding layer (proteins or compounds) and the last two fully connected layers while fixing the RNN, attention and CNN layers of the model. Parameters for those fine-tuned layers were initialized at values in the original models and optimized over the new training data. Batch sizes were set at 2, 10, and 64 for ultra-low (1%), low (5%, 10%), and medium (30%, 50%) training coverage of each set, respectively.

We held out 30% of each labeled generalization set for testing and incrementally made available {1%,5%,10%,30%,50%} of each such set for training.

#### 1.7 Paired t-test for analyzing binding site predictions

To show that the attention values of the binding site are statistically more significant in comparison with the non-binding sites, we performed a one-sided paired t-test. Suppose that there are $n_1$ binding site SSEs and $n_2$ non-binding site ones, we create pairs of samples by pairing each of the binding site SSE with the non-binding site SSE and create $n_1 \times n_2$ such pairs. We assume the null hypothesis that there is no difference between the means of the attention values for binding site SSEs and those for non-binding site SSEs and the alternative hypothesis that the mean of attention values for binding site SSEs are larger than the non-binding site ones. Then we performed the paired t-test as following:

- Calculate the difference between the observations in any pair, $d$.
- Calculate the mean of differences over all $n$ pairs, $\bar{d}$, and their standard deviation $\sigma_d$.
- Calculate the standard error of mean difference by $SE(\bar{d}) = \frac{\sigma_d}{\sqrt{(n)}}$.

- Calculate the t-statistic by $T = \frac{\bar{d}}{SE(\bar{d})}$ under the assumption that this statistic follows a t-distribution with $n-1$ degrees of freedom.
- At the end, calculate the p-value based on the t-distribution.

##### 1.8 Comparing SPS & protein sequence in unsupervised representation learning

We used the same bidirectional+attention seq2seq model for both SPS and amino acid sequences. Since the maximum length of protein sequences in our dataset is 1,500, we used 1,500 two-layer GRUs (almost 10 times that of the SPS seq2seq model with 152 GRUs). For protein sequence data, due to the limit of GPU memory in our facility (one Tesla K80 GPU with 12GB memory), the highest model specification we could try was with the batch size of 8 and the hidden dimension of 64. We trained the protein sequence seq2seq model for 8 days which is twice the limit for training SPS seq2seq model.

##### 1.9 Unified RNN/GCNN-CNN

We compared 1D SMILES representations with the very recently developed graph representations (Gao *et al.*, 2018) for compound protein interactions presented in algorithm 3. In their graphical representation, they used terminology of radius instead of layers. Since they didn't provide their implementation, we implemented their method with their hyper-parameters. We used three layers of GCNN ($R = 3$) and five different convolutional filters instead of one for atoms with different number of neighbors ($H_1^1 \cdots H_R^5$). For example, if an atom has $n$ neighbors then $H^n$ convolutional filter will be used for it in the CNN.

The GCNN described above replaced RNN for compound graphs and RNN was still used for protein sequences. The rest of the unified RNN-CNN stayed the same and led to a unified RNN/GCNN-CNN model.

To be fair, similar to our training process for unified RNN-CNN, we used two phases of training for unified RNN/GCNN-CNN: 1) fixing the protein encoder part and warming-up the rest with learning rate of 0.001; and 2) jointly training all of them together with the lower learning rate of 0.0001.

**Algorithm 3** Graph CNN

**Input:** Molecule graph $G = (V, E)$, radius R, hidden weights $H_1^1 \cdots H_R^5$
**Output:** A vector $\mathbf{r}_a$ for each atom $a$
**Initialize:** Initialize all the $\mathbf{r}_a$
**for** L = 1 to R **do**
  **for** each node a ∈ V **do**
    N = neighbors(a)
    $\mathbf{v} \leftarrow \mathbf{r}_a + \sum_{u \in N} \mathbf{r}_u$
    $\mathbf{r}_a \leftarrow \sigma(\mathbf{v} H_L^{|N|})$
  **end for**
**end for**

### 2 Affinity Prediction Results

#### 2.1 Unsupervised learning for representation pre-training

| | seq2seq | +bucketing | +fw/bw | +attention | +attention+fw/bw |
|---|---|---|---|---|---|
| Number of iterations | 400K | 400K | 400K | 400K | 400K |
| Training error (perplexity) | 7.07 | 7.09 | 2.02 | 1.25 | **1.001** |
| Testing error (perplexity) | 6.5 | 6.91 | 1.84 | 1.13 | **1.0002** |
| Time (h) | 40.2 | 49.52 | 62.93 | 84.75 | 82.52 |

Table S3: Performance comparison among 5 variants of seq2seq for compound representation based on perplexity under the limit of 4-day running time and 400K iterations.

| | seq2seq | +bucketing | +fw/bw | +attention | +attention+fw/bw |
|---|---|---|---|---|---|
| Number of iterations | 400K | 400K | 400K | 153K | 153K |
| Training error (perplexity) | 40.85 | 40.52 | 16.77 | 1.007 | **1.003** |
| Testing error (perplexity) | 41.03 | 43.19 | 19.62 | 1.001 | **1.001** |
| Time (h) | 80.7 | 83.75 | 79.89 | 96 | 96 |

Table S4: Performance comparison among 5 variants of seq2seq for protein representations based on perplexity under the limit of 4-day running time and 400K iterations.

#### 2.2 Supervised shallow learning for representation comparison

| | Baseline representations | | | Novel representations | | |
|---|---|---|---|---|---|---|
| | Ridge | Lasso | RF | Ridge | Lasso | RF |
| Training | 1.23 (0.60) | 1.21 (0.62) | 0.83 (0.84) | 1.26 (0.58) | 1.26 (0.58) | **0.67** (0.91) |
| Testing | 1.24 (0.60) | 1.22 (0.61) | **0.97** (0.78) | 1.27 (0.58) | 1.27 (0.58) | **0.97** (0.78) |
| ER | 1.37 (0.10) | 1.38 (0.10) | 1.35 (0.24) | 1.50 (0.17) | **1.34** (0.14) | 1.48 (0.14) |
| Ion Channel | 1.44 (0.14) | 1.44 (0.14) | **1.38** (0.24) | 1.52 (0.10) | 1.66 (0.10) | 1.46 (0.21) |
| GPCR | 1.27 (0.20) | 1.28 (0.17) | 1.23 (0.30) | 1.44 (0.10) | 1.44 (0.10) | **1.20** (0.19) |
| Tyrosine Kinase | 1.41 (0.41) | 1.44 (0.38) | **1.43** (0.52) | 1.73 (0.20) | 1.77 (0.16) | 1.75 (0.10) |
| Time (core hours) | 2 | 2.22 | 333.2 | 0.22 | 1.42 | 236.21 |
| Memory (GB) | 3.2 | 3.3 | 3.6 | 3.2 | 3.1 | 2.8 |

Table S5: Comparing the baseline and the novel representations based on RMSE (and Pearson correlation coefficient r) of p$K_i$ shallow regression.

| | Baseline representations | | | Novel representations | | |
|---|---|---|---|---|---|---|
| | Ridge | Lasso | RF | Ridge | Lasso | RF |
| Training | 1.04 (0.72) | 1.10 (0.70) | 0.97 (0.78) | 1.17 (0.64) | 1.21 (0.61) | **0.76** (0.88) |
| Testing | 1.33 (0.67) | 1.19 (0.65) | 1.11 (0.70) | 1.24 (0.60) | 1.28 (0.55) | **1.10** (0.70) |
| Ion Channel | **1.53** (0.46) | 1.62 (0.37) | 1.74 (0.30) | 1.81 (0.18) | 1.77 (0.16) | 1.79 (0.09) |
| GPCR | 1.75 (0.07) | 1.74 (0.07) | 1.58 (0.09) | 1.55 (0.10) | **1.53** (0.13) | 1.57 (0.07) |
| Tyrosine Kinase | 1.35 (0.45) | 1.36 (0.45) | **1.32** (0.50) | 1.56 (0.18) | 1.59 (0.13) | 1.46 (0.31) |
| Time (core hours) | 0.1 | 0.16 | 4.78 | 0.01 | 0.06 | 10.34 |
| Memory (Gb) | 0.4 | 0.4 | 0.4 | 0.4 | 0.4 | 0.4 |

Table S6: Comparing the baseline and the novel representations based on RMSE (and Pearson correlation coefficient $r$) of p$K_d$ shallow regression.

| | Baseline representations | | | Novel representations | | |
|---|---|---|---|---|---|---|
| | Ridge | Lasso | RF | Ridge | Lasso | RF |
| Training | 1.02 (0.72) | 0.98 (0.74) | 0.74 (0.86) | 0.92 (0.78) | 0.97 (0.74) | **0.64** (0.90) |
| Testing | 1.04 (0.70) | 1.02 (0.72) | 0.90 (0.79) | 0.93 (0.77) | 0.98 (0.74) | **0.80** (0.84) |
| ER | 1.55 (0.20) | 1.60 (0.17) | 1.71 (0.17) | 1.72 (0.14) | 1.55 (0.17) | **1.51** (0.13) |
| Ion Channel | 1.25 (0.44) | 1.26 (0.43) | 1.21 (0.48) | 1.30 (0.40) | 1.39 (0.30) | **1.20** (0.33) |
| GPCR | 1.34 (0.20) | 1.38 (0.14) | 1.39 (0.14) | 1.31 (0.24) | 1.34 (0.22) | **1.31** (0.24) |
| Tyrosine Kinase | 1.25 (0.09) | 1.30 (0.08) | 1.21 (0.09) | 1.33 (0.25) | 1.63 (0.26) | **1.07** (0.10) |
| Time (core hours) | 0.34 | 0.42 | 38.46 | 0.05 | 0.26 | 31.67 |
| Memory (GB) | 1 | 1 | 1.1 | 0.9 | 0.9 | 0.8 |

Table S7: Comparing the baseline and the novel representations based on RMSE (and Pearson correlation coefficient $r$) of $pEC_{50}$ shallow regression.

#### 2.3 Supervised deep learning for p$K_i$ prediction

| | RF | Unified RNN-CNN Models (separate attention) | | | Unified RNN-CNN Models (joint attention) | | |
|---|---|---|---|---|---|---|---|
| | | single | parameter ensemble | parameter+NN ensemble | single | parameter ensemble | parameter+NN ensemble |
| Training | 0.67 (0.91) | 0.44 (0.95) | 0.42 (0.95) | 0.41 (0.96) | 0.48 (0.94) | 0.44 (0.95) | **0.40** (0.96) |
| Testing | 0.97 (0.78) | 0.84 (0.84) | 0.83 (0.84) | **0.79** (0.86) | 0.91 (0.81) | 0.88 (0.82) | 0.80 (0.85) |
| ER | **1.48** (0.14) | 1.76 (0.17) | 1.74 (0.17) | 1.62 (0.07) | 1.76 (0.09) | 1.78 (0.07) | 1.63 (0.10) |
| Ion Channel | 1.46 (0.21) | 1.50 (0.21) | 1.49 (0.20) | **1.41** (0.29) | 1.79 (0.23) | 1.78 (0.24) | 1.51 (0.32) |
| GPCR | **1.20** (0.19) | 1.35 (0.28) | 1.33 (0.29) | 1.26 (0.36) | 1.50 (0.21) | 1.48 (0.23) | 1.35 (0.28) |
| Tyrosine Kinase | **1.75** (0.10) | 1.83 (0.27) | 1.81 (0.27) | 1.85 (0.25) | 2.10 (0.16) | 2.08 (0.15) | 1.95 (0.17) |

Table S8: Under novel representations learned from seq2seq, comparing random forest and variants of separate RNN-CNN and unified RNN-CNN models based on RMSE (and Pearson correlation coefficient $r$) for p$K_i$ prediction.

#### 2.4 Generalization sets v.s. training and testing sets

##### 2.4.1 Distributions in compound SMILES and protein SPS

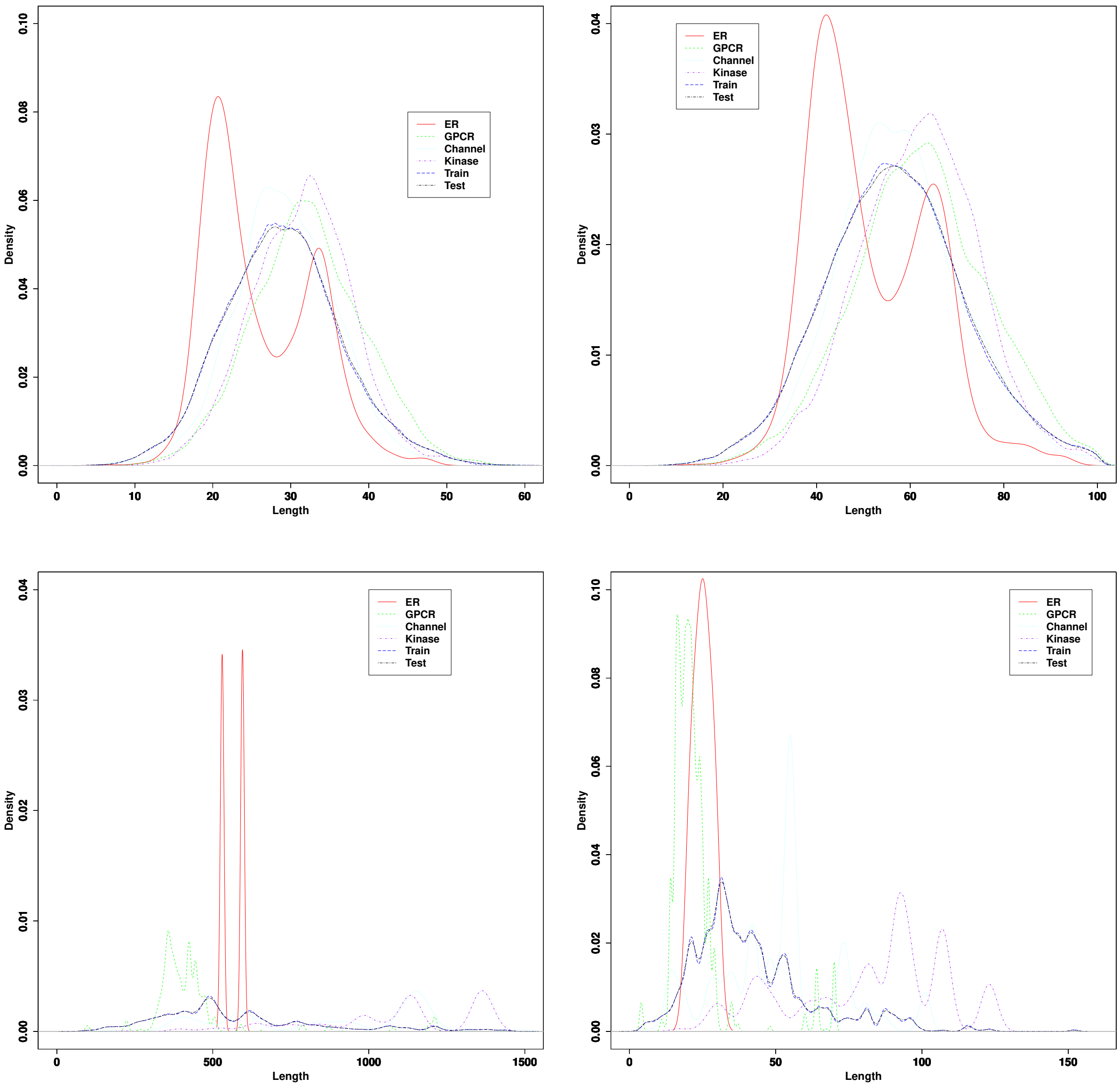


Figure S3: Comparing marginal distributions of various sets of compounds in size, i.e., the number of atoms (top left); compounds in SMILES length (top right); protein targets in the amino-acid length (bottom left); and protein targets in the SPS string length (bottom right).

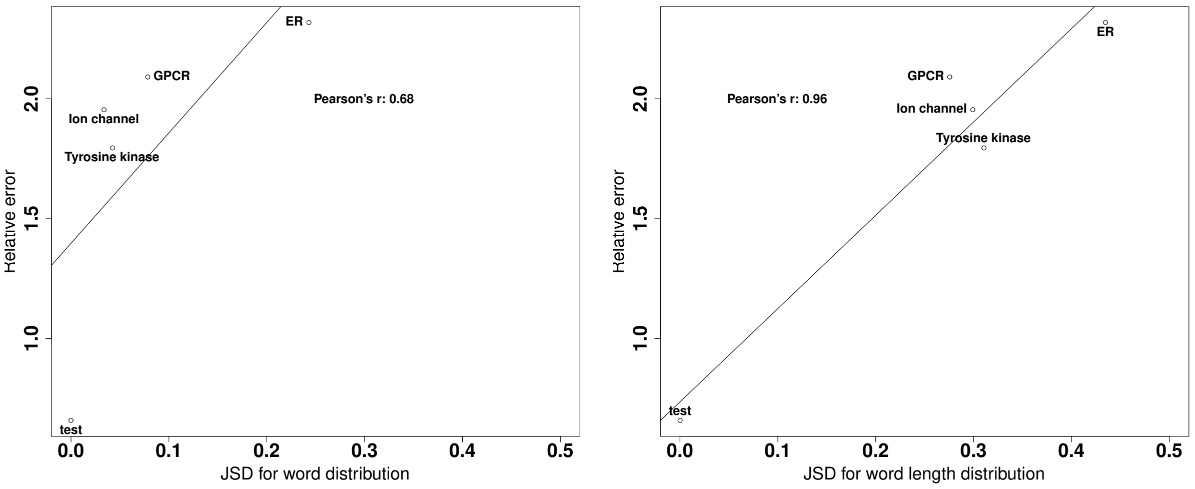


Figure S4: Relative errors to the training set (y axis) versus Jensen-Shannon distances from the training-set protein SPS letter distribution (x axis: left) or SPS length distribution (x axis: right) for various sets of protein targets.

##### 2.4.2 Predicted v.s. measured $\mathbf{pIC_{50}}$

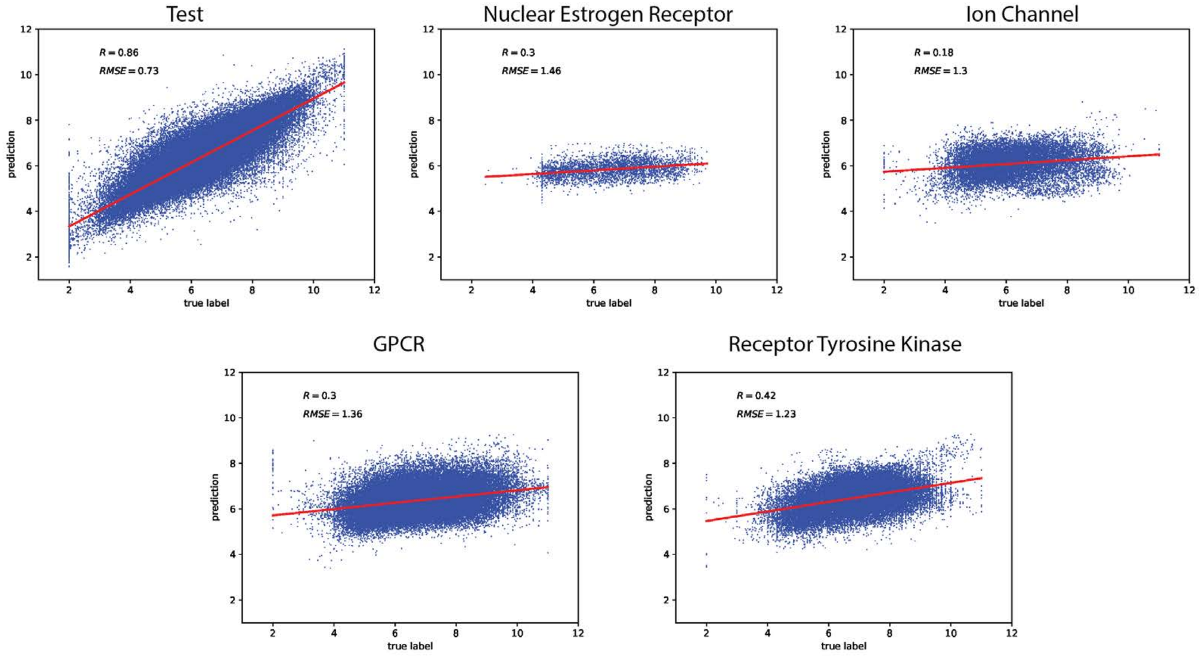


Figure S5: Comparing predictions vs real labels for test and generalization tests for the unified RNN-CNN model (separate attention).

### 3 Interpreting Affinity and Specificity Predictions

#### 3.1 Comparing binding site prediction between separate attention and joint attention

| | | Number of SSEs | | Top 10% (4) SSEs predicted as binding site by sep. attn. | | | | Top 10% (4) SSEs predicted as binding site by joint attn. | | | |
|---|---|---|---|---|---|---|---|---|---|---|---|
| Target–Drug | PDB ID | total | binding site | # of TP | Enrichment | Highest rank | P value | # of TP | Enrichment | Highest rank | P value |
| Human COX2–rofecoxib | 5KIR | 40 | 6 | 2 | 2.22 | 1 | 1.18e-9 | 1 | 1.68 | 4 | 0.0107 |
| Human PTP1B–OBA | 1C85 | 34 | 5 | 0 | 0 | 20 | 1 | 1 | 1.7 | 1 | 1.12e-10 |
| Human factor Xa–DX9065 | 1FAX | 31 | 4 | 0 | 0 | 7 | 0.898 | 3 | 5.81 | 2 | 2.2e-16 |

Table S9: Interpreting deep learning models: predicting binding sites based on joint attentions. The binding site here is defined as SSEs making direct contacts with compounds (according to the LIGPLOT service from PDBsum).

| | | Number of SSEs | | Top 10% (4) SSEs predicted as binding site by joint attn. | | | |
|---|---|---|---|---|---|---|---|
| Target–Drug | PDB ID | total | binding site | # of TP | Enrichment | Highest rank | P value |
| Human COX2–rofecoxib | 5KIR | 40 | 9 | 1 | 1.11 | 4 | 1.3e-1 |
| Human PTP1B–OBA | 1C85 | 34 | 6 | 2 | 3.77 | 1 | <2.2e-16 |
| Human factor Xa–DX9065 | 1FAX | 31 | 6 | 4 | 5.16 | 1 | <2.2e-16 |

Table S10: Interpreting deep learning models: predicting binding sites based on joint attentions. The binding site here is defined as SSEs falling within 5Å from compound heavy atoms.

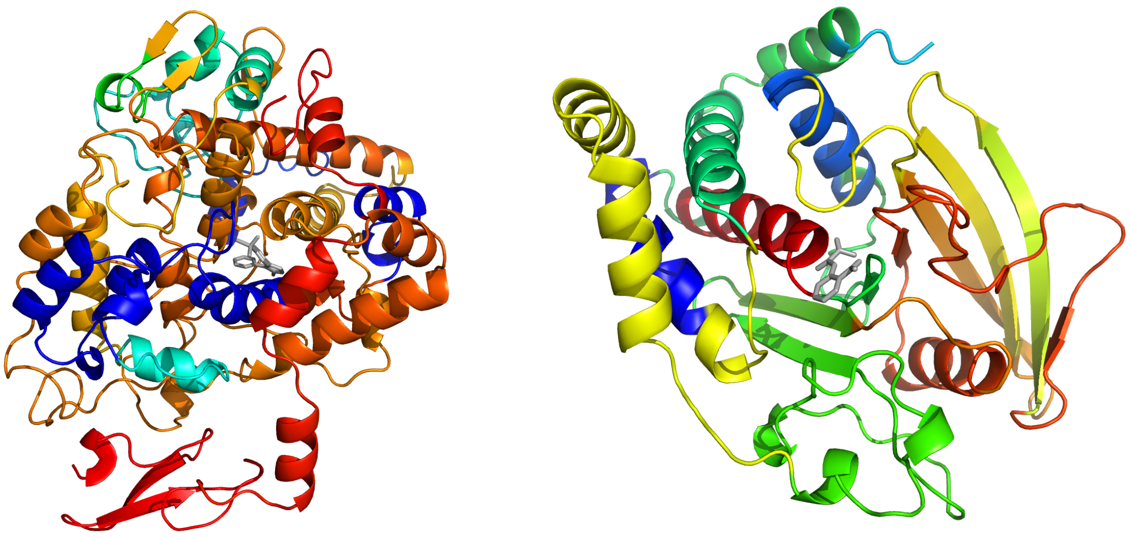

Figure S6: Protein binding site interpretation for more protein-compound pairs: (Left) COX2-rofecoxib (PDB ID: 5KIR) and (Right) PTP1B-OBA (PDB ID:1C85). Proteins are shown in cartoons and color-coded with attention scores $\beta_i$. Compounds are shown in black sticks.

##### 3.2 Attention scores on compounds

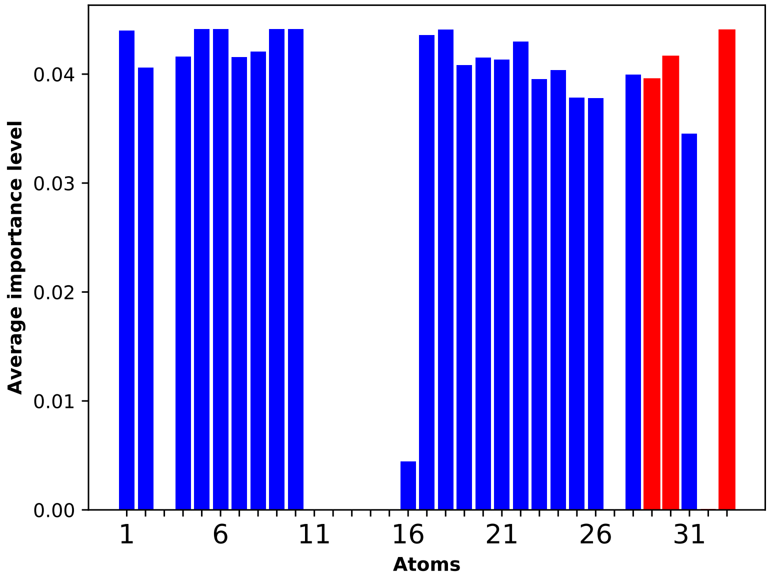


Figure S7: Max-marginalized attention scores $\beta_j$'s for compound DX-9065a interacting with factor Xa.

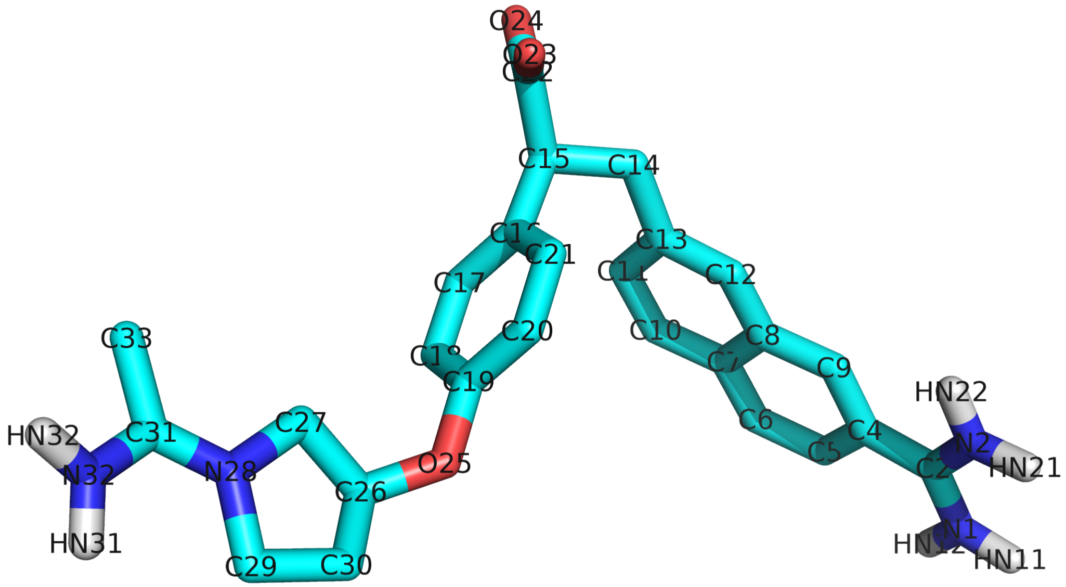


Figure S8: The chemical structure and atom names for compound DX-9065a, a selective ligand for factor Xa.

| index | atom name | index | atom name |
|---|---|---|---|
| 1 | C33 | 18 | C13 |
| 2 | C31 | 19 | C12 |
| 3 | N32 | 20 | C8 |
| 4 | N28 | 21 | C7 |
| 5 | C29 | 22 | C10 |
| 6 | C30 | 23 | C11 |
| 7 | C26 | 24 | C6 |
| 8 | C27 | 25 | C5 |
| 9 | O25 | 26 | C4 |
| 10 | C19 | 27 | C9 |
| 11 | C18 | 28 | C2 |
| 12 | C17 | 29 | N1 |
| 13 | C16 | 30 | N2 |
| 14 | C21 | 31 | C22 |
| 15 | C20 | 32 | O23 |
| 16 | C15 | 33 | O24 |
| 17 | C14 | - | - |

Table S11: Correspondence between atom indices of compound DX-9065a in Fig. S7 and atom names in Fig. S8.

#### 3.3 Selectivity origin prediction

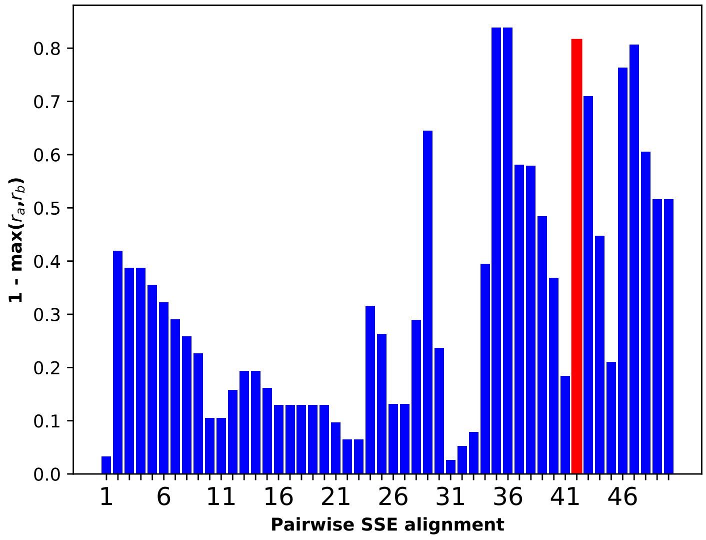


Figure S9: Interpreting joint attention in our unified RNN-CNN model for factor Xa specificity. Pairwise alignment of amino-acid sequences of factor Xa and thrombin decomposed both sequences into 50 segments. These segments are scored by an alternative measure: one less the maximum between the corrected attention rank ratios for the two compound-protein interactions. The ground truth of specificity origin is in red.

| Segment (paired SSE) index | Thrombin SSE index | Factor Xa SSE index | Segment (paired SSE) index | Thrombin SSE index | Factor Xa SSE index |
|---|---|---|---|---|---|
| 1 | 1 | 1 | 26 | 17 | 16 |
| 2 | 1 | 2 | 27 | 17 | 17 |
| 3 | 2 | 3 | 28 | 18 | 17 |
| 4 | 3 | 3 | 29 | 19 | 17 |
| 5 | 3 | 4 | 30 | 20 | 17 |
| 6 | 4 | 5 | 31 | 21 | 18 |
| 7 | 4 | 6 | 32 | 22 | 19 |
| 8 | 4 | 7 | 33 | 23 | 20 |
| 9 | 4 | 8 | 34 | 24 | 21 |
| 10 | 5 | 8 | 35 | 25 | 21 |
| 11 | 5 | 9 | 36 | 26 | 21 |
| 12 | 6 | 9 | 37 | 27 | 22 |
| 13 | 7 | 9 | 38 | 28 | 23 |
| 14 | 8 | 9 | 39 | 28 | 24 |
| 15 | 8 | 10 | 40 | 29 | 24 |
| 16 | 8 | 11 | 41 | 30 | 25 |
| 17 | 9 | 11 | 42 | 31 | 26 |
| 18 | 10 | 11 | 43 | 32 | 27 |
| 19 | 11 | 11 | 44 | 33 | 27 |
| 20 | 12 | 11 | 45 | 34 | 27 |
| 21 | 13 | 12 | 46 | 35 | 28 |
| 22 | 13 | 13 | 47 | 36 | 29 |
| 23 | 14 | 13 | 48 | 37 | 30 |
| 24 | 15 | 14 | 49 | 37 | 31 |
| 25 | 16 | 15 | 50 | 38 | 31 |

Table S12: Correspondence between segment indices in Fig. S9 (as well as Fig. 4 in Main Text) and the paired/aligned indices of target (factor Xa) and off-target (thrombin) SSEs.

### References


Abadi, M., Barham, P., Chen, J., Chen, Z., Davis, A., Dean, J., Devin, M., Ghemawat, S., Irving, G., Isard, M., *et al.* (2016). Tensorflow: A system for large-scale machine learning. In *OSDI*, volume 16, pages 265–283.

Bahdanau, D., Cho, K., and Bengio, Y. (2014). Neural machine translation by jointly learning to align and translate. *arXiv preprint arXiv:1409.0473*.

Britz, D., Goldie, A., Luong, T., and Le, Q. (2017). Massive Exploration of Neural Machine Translation Architectures. *ArXiv e-prints*.

Gao, K. Y., Fokoue, A., Luo, H., Iyengar, A., Dey, S., and Zhang, P. (2018). Interpretable drug target prediction using deep neural representation. In *IJCAI*, pages 3371–3377.

Glorot, X. and Bengio, Y. (2010). Understanding the difficulty of training deep feedforward neural networks. In *Proceedings of the thirteenth international conference on artificial intelligence and statistics*, pages 249–256.

Khomenko, V., Shyshkov, O., Radyvonenko, O., and Bokhan, K. (2016). Accelerating recurrent neural network training using sequence bucketing and multi-gpu data parallelization. In *Data Stream Mining & Processing (DSMP), IEEE First International Conference on*, pages 100–103. IEEE.

Kim, S., Thiessen, P. A., Bolton, E. E., Chen, J., Fu, G., Gindulyte, A., Han, L., He, J., He, S., Shoemaker, B. A., Wang, J., Yu, B., Zhang, J., and Bryant, S. H. (2016). PubChem Substance and Compound databases. *Nucleic Acids Res.*, **44**(D1), D1202–1213.

Lawrence, S., Giles, C. L., Tsoi, A. C., and Back, A. D. (1997). Face recognition: A convolutional neural-network approach. *IEEE transactions on neural networks*, **8**(1), 98–113.

Liu, T., Lin, Y., Wen, X., Jorissen, R. N., and Gilson, M. K. (2006). Bindingdb: a web-accessible database of experimentally determined protein–ligand binding affinities. *Nucleic acids research*, **35**(suppl_1), D198–D201.

Lu, J., Yang, J., Batra, D., and Parikh, D. (2016). Hierarchical question-image co-attention for visual question answering. In *Advances In Neural Information Processing Systems*, pages 289–297.

Maas, A. L., Hannun, A. Y., and Ng, A. Y. (2013). Rectifier nonlinearities improve neural network acoustic models. In *Proc. icml*, volume 30, page 3.

Pedregosa, F., Varoquaux, G., Gramfort, A., Michel, V., Thirion, B., Grisel, O., Blondel, M., Prettenhofer, P., Weiss, R., Dubourg, V., Vanderplas, J., Passos, A., Cournapeau, D., Brucher, M., Perrot, M., and Duchesnay, E. (2011). Scikit-learn: Machine learning in Python. *Journal of Machine Learning Research*, **12**, 2825–2830.

Schuster, M. and Paliwal, K. K. (1997). Bidirectional recurrent neural networks. *IEEE Transactions on Signal Processing*, **45**(11), 2673–2681.

Tang, Y. (2016). Tf. learn: Tensorflow's high-level module for distributed machine learning. *arXiv preprint arXiv:1612.04251*.
